# Quantification of proteins, protein complexes and mRNA in single cells by proximity-sequencing

**DOI:** 10.1101/2020.05.15.098780

**Authors:** Luke Vistain, Hoang Van Phan, Christian Jordi, Mengjie Chen, Sai T. Reddy, Savaş Tay

**Affiliations:** Pritzker School of Molecular Engineering, The University of Chicago, Chicago, IL 60637, USA; Institute for Genomics and Systems Biology, The University of Chicago, Chicago, IL 60637, USA; Department of Biosystems Science and Engineering, ETH Zurich, Basel, 4058, Switzerland; Section of Genetic Medicine, Department of Medicine, The University of Chicago, Chicago, IL 60637, USA; Department Human Genetics, The University of Chicago, Chicago, IL 60637, USA

## Abstract

Multiplexed analysis of single-cells enables accurate modeling of cellular behaviors, classification of new cell types, and characterization of their functional states. Here we present proximity-sequencing (Prox-seq), a method for simultaneous measurement of an individual cell’s proteins, protein complexes and mRNA. Prox-seq utilizes deep sequencing and barcoded proximity assays to measure proteins and their complexes from all pairwise combinations of targeted proteins, in thousands of single-cells. The number of measured protein complexes scales quadratically with the number of targeted proteins, providing unparalleled multiplexing capacity. We developed a high-throughput experimental and computational pipeline and demonstrated the potential of Prox-Seq for multi-omic analysis with a panel of 13 barcoded proximity probes, enabling the measurement of 91 protein complexes, along with thousands of mRNA molecules in single T-cells and B-cells. Prox-seq provides access to an untapped yet powerful measurement modality for single-cell phenotyping and can discover new protein interactions in signaling and drug studies.

---

Singe-cell measurements have expanded our understanding of many aspects of cellular function, such as enabling identification of rare cell subsets, tracking transient cellular states, and incorporating noise and variability into our understanding of cellular phenotypes^1–3^. These phenotypes are emergent properties of both the biomolecules in the cell and the interaction between these molecules. Therefore, the ability to measure individual proteins and their complexes on a single-cell level would be among the most informative measurements for understanding cellular function. Despite the apparent value, there are major hurdles in making highly multiplexed measurements of single-cell proteins and their complexes. Combining high-dimensional multiplexing with single-cell resolution demands a method that can encode a large number of outputs, as each measurement must enable identification of both the protein of interest and other proteins in its proximity, along with their transcripts.

Motivated by this unmet need in multi-omic analysis in single cells, we developed an end-to-end experimental and computational pipeline called Prox-Seq (Fig. 1a), for simultaneous measurement of proteins, their complexes and mRNA in up to hundreds of thousands of individual cells. Our method integrates Proximity Ligation Assay (PLA)^4^ and deep sequencing for detection of proteins and their complexes. PLA uses pairs of DNA-conjugated antibodies (PLA probes) designed in such a way that, upon being in proximity (both antibodies bind to epitopes that are spatially proximal), the DNA oligomers can ligate. This method has proven sensitive enough to measure proteins from single cells and the abundance of protein can be measured via DNA sequencing^5–7^. In this way, one can encode both the identity of a protein and its proximity partner into the formation of a unique pair of DNA barcodes, which can then leverage the multiplexing capacity of a deep sequencing readout.

**Fig. 1:**
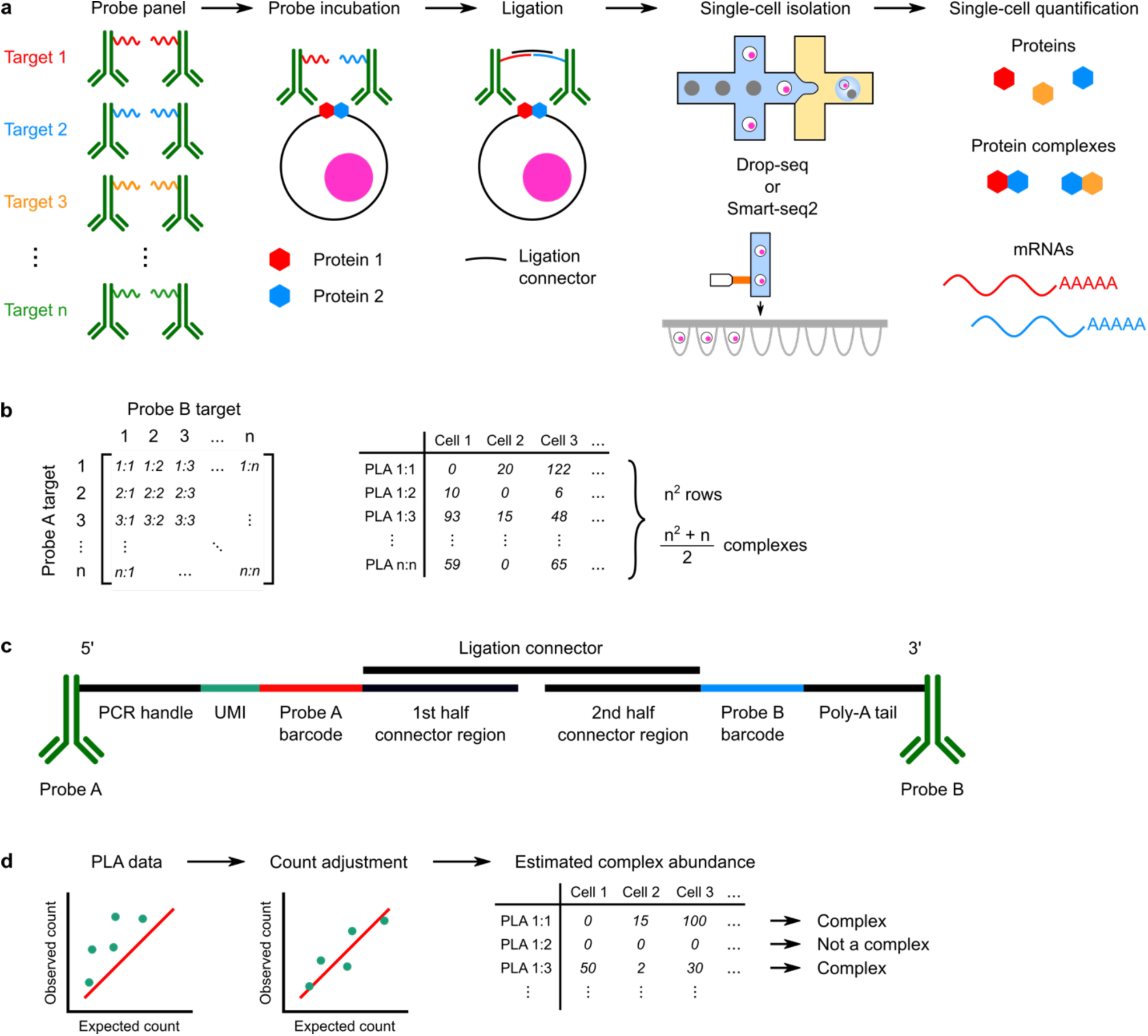
Overview of Prox-seq for joint proteomic and transcriptomic analysis of single cells. **a**, Prox-seq workflow: cells are stained with barcoded DNA-conjugated antibodies (proximity probes), ligated, and are processed using a Drop-seq or Smart-seq2 protocol. Sequencing readout simultaneously reveals protein, protein-complex, and mRNA abundance. **b**, The protein output of Prox-seq is a set of n^2^ PLA products, representing up to (n^2^+n)/2 possible protein dimers and interactions. **c**, Design of the Prox-seq probes and the DNA oligonucleotide (oligomer) on the antibodies. Probes A and probes B are conjugated with the oligomers at the 5’ and 3’ end, respectively. **d**, Data analysis workflow detecting protein complexes and estimating their abundances. The count of each PLA product is compared to the expected value in the absence of stable protein interactions. Additional counts above the expected value indicate the presence of a protein complex.

Prox-seq expands on the state of the art by enabling measurements of protein proximity from all pairwise combinations of targeted proteins^8,9^ from hundreds of thousands of cells and has immense potential for highly-multiplexed proteomic analysis. The number of possible protein complexes scales quadratically with the number of targeted proteins (Fig. 1b). For example, a panel of 100 antibodies can measure more than 5000 unique pairwise complexes, well beyond the measurement capacity of current single-cell proteomic methods and approaching the multiplexing capacity of single-cell RNA sequencing. In addition to measuring proteins and their complexes, our approach readily enables the measurement of the transcriptome of each cell via RNA sequencing, providing a uniquely multiplexed mode of single cell measurements.

For each protein target in our panel we generate a pair of Prox-seq PLA probes (probe A and B) with single-stranded oligonucleotides (oligomers) (Fig. 1c). The oligomers are designed such that each member of group A can ligate with any member of group B. After antibody binding and ligation, cells can be processed through commonly used single-cell sequencing (scRNA-seq) methods, such as Drop-seq or Smart-seq2^10,11^, to retrieve both PLA products and mRNAs. Thus, our method can be seamlessly integrated in existing research workflows. The oligomers used to form PLA products include several features that facilitate integration into these single-cell sequencing methods (Fig. 1c). The complete PLA product includes a unique molecular identifier (UMI) region for read count normalization, two sequence barcodes to indicate the identity of the A and B antibodies, a poly-A tail for capture, and a primer binding site for PCR. The connector regions enable proximity ligation, and only ligated products can be PCR amplified. Finally, the data are processed to identify which PLA products reflect the existence of a protein dimer (Fig. 1d).

We first sought to show that PLA products can be measured using single-cell sequencing, and that the PLA data display cell type-specific differences. Eleven protein targets were selected corresponding to various markers of T-cell and B-cell lymphocytes (Supplementary table 1). PLA probes were made for these targets along with two isotype controls. This panel was applied to a mixture of T-cells (Jurkat cells) and B-cells (Raji cells). Individual cells were analyzed using the Drop-seq method^11^ (Supplementary Fig 1a). Following Prox-seq, we found that cells could be accurately clustered using either mRNA, protein abundance, or PLA products (Fig. 2a-c). Further, clustering by mRNA or protein identified the same cell types (Fig. 2b, d). The protein abundance of a target is estimated from the total number of times the target’s DNA barcode is detected, from either PLA probe A or B. Similarly, cells could be clustered using all 169 PLA products, which reflect proximity information rather than strict protein abundance (Fig. 2c). Regardless of the data used to cluster the cells, Prox-seq displayed good concordance between the level of gene expression and protein for a given cluster (Fig. 2e, Supplementary Fig. 2). We also found that CD3:CD147 and CD3:CD3 were the two most significantly upregulated PLA products in the Jurkat cluster. On the other hand, ICAM1:HLA-DR and HLA-DR:HLA-DR were the two most significantly upregulated ones in the Raji cluster (Fig. 2f). Finally, we analyzed the same antibody panel with a Smart-seq2 method and showed that the rank order of protein abundance level for each cell line was comparable between Prox-seq data and flow cytometry, indicating good agreement between the two methods (Fig. 2g, Supplementary Figs. 1b, 3, 4). This demonstrates that Prox-seq recapitulates the features of other similar assays such as REAP-seq and CITE-seq for protein quantification^8,9^.

**Fig. 2:**
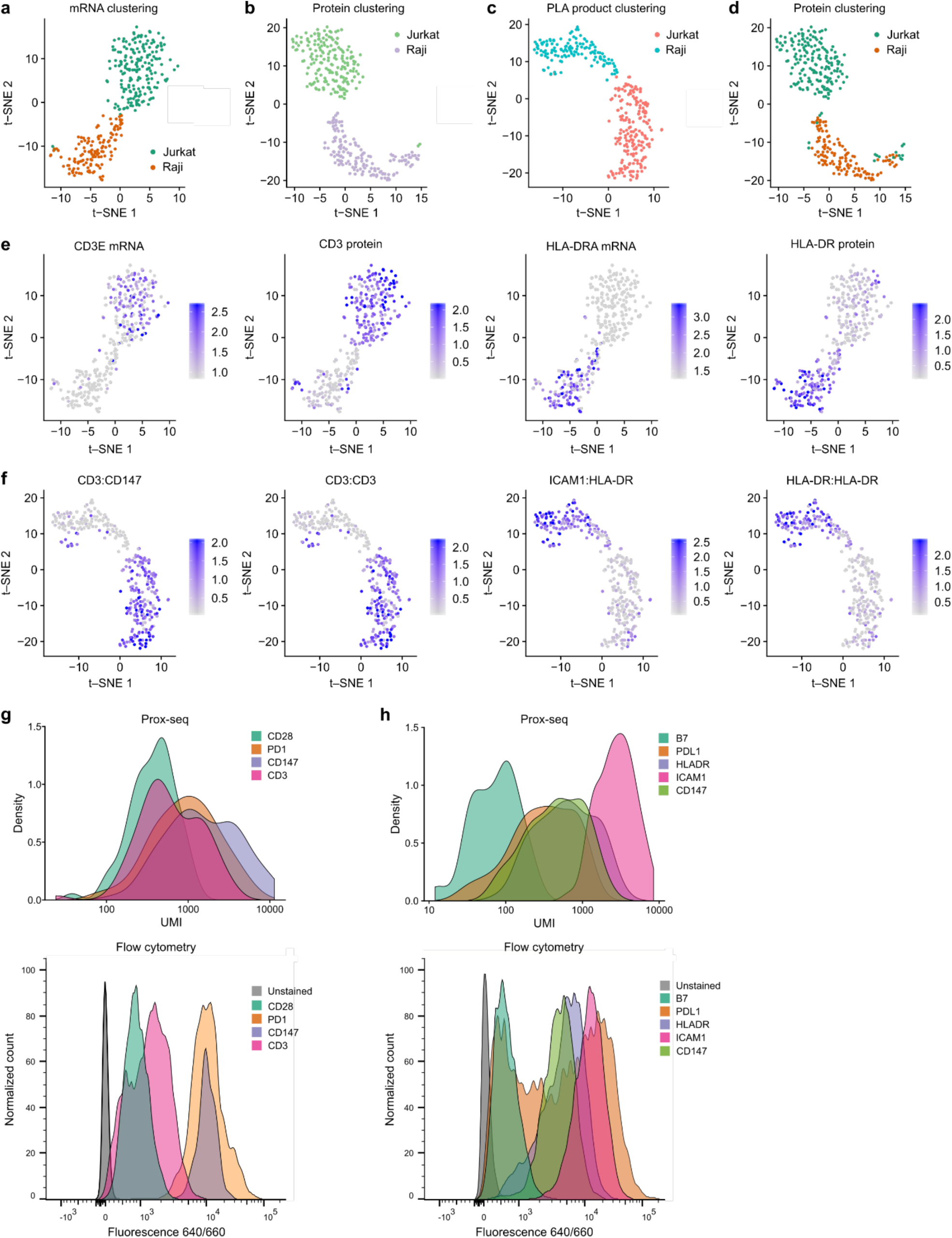
Prox-seq measurements identify cell types and accurately reflect protein abundance in single-cells. **a-c**, We measured a T-cell line (Jurkat) and B-cell line (Raji) with our method. Prox-seq measures proteins, protein complexes and mRNA simultaneously in single-cells. t-SNE plots showing cell clustering using (a) mRNA data, (b) protein abundance data, and (c) PLA data. **d**, The same t-SNE plot as in (b), but with clusters labeled using mRNA data. **e**, Concordance between protein and mRNA levels for the CD3D gene, CD3 protein, HLA-DRA gene, and HLA-DR protein. **f**, The most significant cluster markers are CD3:CD147, CD3:CD3, ICAM1:HLA-DR, and HLA-DR:HLA-DR. **g**, Comparison of T cell protein abundance between Prox-seq data (top) and flow cytometry data (bottom) for the same cells. **h**, Comparison of B cell protein abundance between Prox-seq data (top) and flow cytometry data (bottom) for B-cells shows good agreement in rank order of individual proteins. (a-f) Data was generated using the Drop-seq method. (g-h) Data was generated using the Smart-seq2 method.

A unique feature of Prox-seq is that it can reveal pairwise protein interaction information for each of the targeted proteins. To demonstrate this capability we again treated Jurkat and Raji cells with our panel of 13 Prox-seq probes, and analyzed the PLA products using a modified Smart-seq2 protocol^10^ (Supplementary Methods). This panel allows us to measure up to 91 potential pairwise protein complexes (Fig. 1b). The generation of a PLA product in Prox-seq indicates that two proteins were close enough to ligate the two 60-base long DNA strands. To verify that PLA protein-complex products are not occurring simply by chance (i.e., random ligation of two non-interacting proteins, Supplementary Fig. 5) and that the two proteins are part of a stable protein complex^12^, we performed statistical modeling and compared them to our data. In the absence of stable protein complexes, the expected count of all PLA products can be calculated from the abundance of protein targets (Supplementary Methods). We verified this using a model based on the Poisson point process to simulate Prox-seq PLA data in the presence and absence of protein complex information (Supplementary Fig. 6, Supplementary Methods). Our experimental data showed that for several PLA products, the observed PLA count was significantly higher than what should be expected by chance (Supplementary Figs. 7, 8). Linear regression showed that the interaction effect was significant in explaining the observed PLA abundance, thus showing stable complex formation (approximately 40% and 17% of PLA products in Jurkat and Raji cells, respectively, had a P value lower than 0.05 (two-sided likelihood ratio test), Supplementary Fig. 9, Supplementary Methods).

The protein complexes measured by Prox-seq can be analyzed through simple visual inspection (Fig. 3a-b). However, we also sought to identify which PLA products measure protein complexes, and how abundant the complexes were. To accomplish this, we developed a Protein Complex Estimation Algorithm (Supplementary Methods) and tested it on simulated data. We found that we were able to detect true protein complexes (Supplementary Fig. 10). A more detailed discussion on the algorithm is available in the Supplementary Methods. Next, we applied this algorithm to our real data set and found that four PLA products were calculated to have more than 50% of their signal being attributed to dimers: CD3 and CD28 homodimers in Jurkat cells, and PDL1 and HLA-DR homodimers in Raji cells (Fig. 3c-f). These four homodimers also have the highest estimated abundance (P<10^−6^ compared to negative control PLA products (two-sided Welch’s t-test), Supplementary Fig. 11). It is important to note that, apart from CD147 and the isotype controls, all antibodies in our panel are monoclonal. This enables detection of both heterodimers and homodimers. The CD3 homodimer is noteworthy because it serves as a positive control in our panel. The antibody targets the CD3ε protein, two of which are part of the T-cell receptor complex^13^. Furthermore, CD28 is known to form a stable homodimer on the cell surface through a disulfide bridge^14^. While it is unclear from previous studies if PDL1 forms a homodimer on the cell surface, all known crystal structures of PDL1 feature a homodimer^15^, similar to what we found in our measurements. HLA-DR is thought to exist in an equilibrium between monomers and homodimers on the cell surface, with homodimers and oligomers being the functional unit for T-cell receptor engagement^16,17^. Therefore, the Protein Complex Estimation Algorithm was able to correctly identify the presence of four known protein complexes. However, B7 and ICAM1 are both thought to undergo some degree of homodimerization^18,19^. ICAM1 does indeed have the highest number of UMI’s attributed to complexes but, due to its very high expression level, that only represents a small fraction of total reads in our data (Fig. 3d, Supplementary Data). The absence of B7 homodimers raises the possibility that the monoclonal antibody in this panel is unable to bind to the dimerized form.

**Fig 3:**
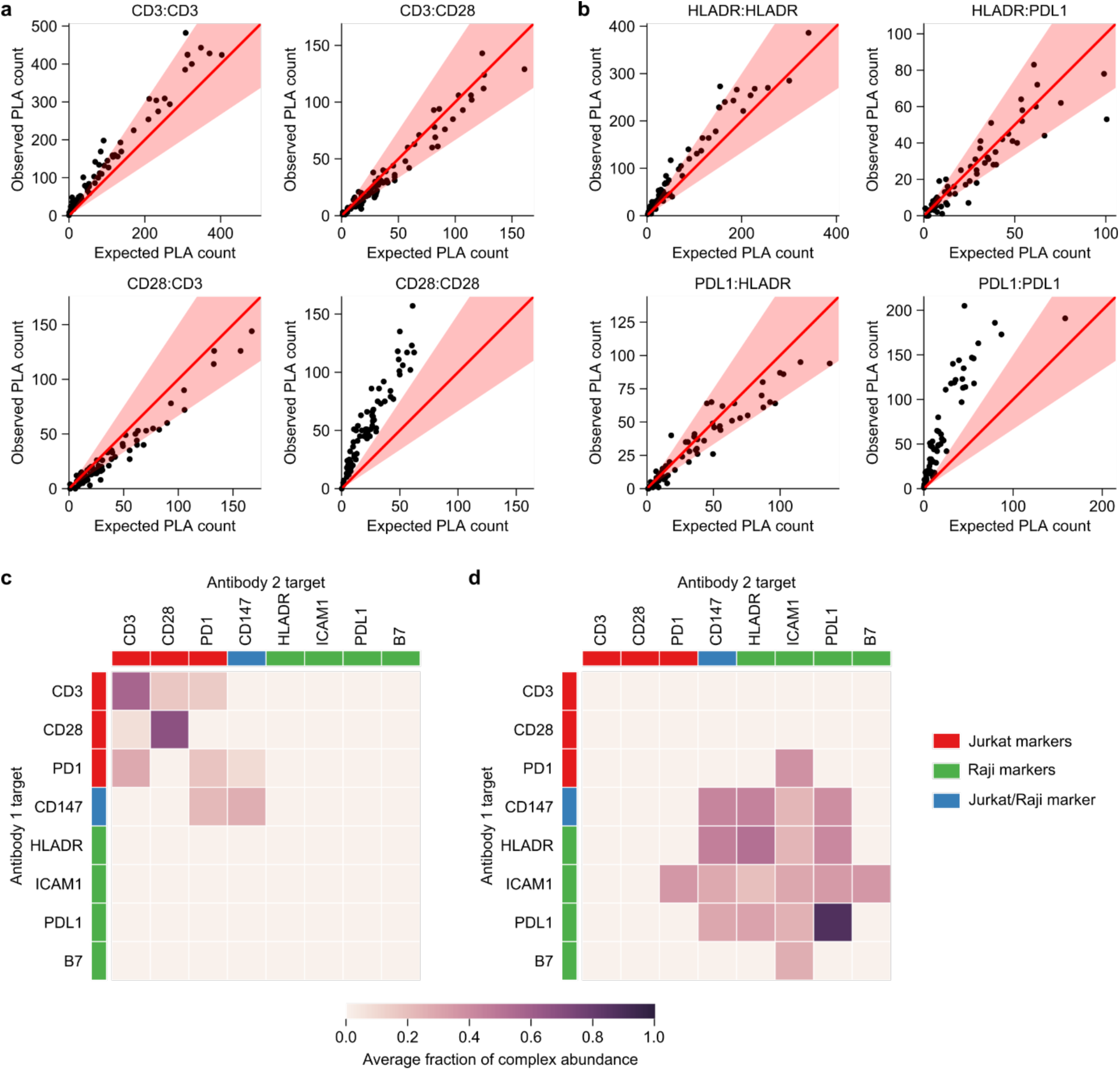
Quantification of protein complexes in single-cells using Prox-seq. We measured the complexes among cell-specific marker proteins, including CD3, CD28, HLADR, and PDL1. **a**, Scatter plots showing observed count against the expected count of 4 example PLA protein-complex products in T-cells (Jurkats). **b**, Scatter plots showing observed count against the expected count of 4 example PLA protein-complex products in B-cells (Rajis). (a, b) The red solid lines indicate the line y=x, and the red regions indicate 1.5-fold change from the expected PLA count. **c**,**d**, Heatmaps showing the estimated protein complex abundance as a fraction of observed PLA count in single (c) Jurkat and (d) Raji cells.

In summary, we demonstrated the ability to make simultaneous protein, protein-complex and mRNA measurements in single-cells, which will be a valuable tool for understanding cellular function, and for classification of cell types and states. Signaling, differentiation, development and decision making in cells is driven in large part by changes in the arrangement of protein molecules. A broader insight into these arrangements can assist in the aspirational goal of constructing models which more accurately predict cellular behavior from knowledge of their constituent parts. We show that this technology is compatible with commonly used single-cell sequencing methods and can be used to identify cell types. Most importantly, Prox-seq can identify members of pairwise protein complexes, providing an entirely new functionality to single-cell sequencing.

Recent advances in single-cell sequencing technology have enabled comprehensive characterization of the transcriptome, genome and epigenome at the single-cell level^11,20,21^. Several methods have expanded these approaches to incorporate antibody-based protein measurements ^8,9,22^. Prox-seq can recapitulate the functionality of these methods in addition to measuring protein complexes, and provides quadratically-scaled multiplexing capability to greatly advance single-cell proteomic analysis. Currently, in order to make highly multiplexed measurements of protein complexes, one is limited to bulk samples. These measurements are most often achieved using mass spectrometry or yeast-two-hybrid assays^23–25^. Conversely, current single-cell measurements are typically restricted to optical techniques, which are limited in their multiplexing capacity due to spectral overlapping, and the compatibility of fluorophore pairs^26,27^. Prox-seq bridges this divide by providing both multiplexing and single-cell resolution. Furthermore, Prox-seq incorporates single-cell RNA sequencing, thereby providing multiple single-cell data types simultaneously. Finally, Prox-seq can be expanded to other modes of analysis by simply changing the antibodies. For example, one can probe post-translation modifications and interactions between proteins and histone modifications by using the appropriate antibody. Therefore, Prox-seq can provide greatly enhanced multi-omic analysis capability in single cells.

## Author contributions

L.V., H.V.P., C.J., S.T.R., and S.T. conceived of and designed the project. L.V., H.V.P., and C.J. performed all experiments, designed the PLA probes’ components, and analyzed sequencing data. H.V.P., and M.C. performed statistical analysis. L.V., H.V.P., and S.T. wrote the manuscript. S.T. supervised the project. All authors reviewed the manuscript.

## Acknowledgements

We thank Jun Huang for his generous gift of the cell line used in this study. We thank Anindita Basu and Heather Eckart for their advice on Drop-seq protocol. We acknowledge both the University of Chicago Genomics Facility, The University of Chicago Cytometry and Antibody Technology facility, and The University of Chicago Research Computing Center for their services. We thank Ulf Landegren for assistance with PLA. This work was supported by the NIH R01 grant GM127527 and Paul G. Allen Distinguished Investigator Award to S.T.

## Methods

### Prox-seq PLA probe preparation

Antibodies were DNA-conjugated using previously published methods^28^. Briefly, 9µL antibody solutions in phosphate buffered saline (PBS) at 1-2 mg/ml were combined with 1 µL 2mM dibenzocyclooctyne-PEG4-*N*-hydroxysuccinimidyl ester (Sigma, Catalog # 764019) in dimethylsulfoxide (DMSO, Sigma Aldrich) and reacted on ice for 1-2 hours. Mouse isotype controls IgG1, IgG2a, and IgG2b were concentrated and buffer exchanged prior to conjugation using a 50K molecular weight cutoff (MWCO) concentrator (EMD Millipore). After incubation the DBCO-conjugated antibodies were purified using a 7K MWCO Zeba desalting column (Life Technologies) and the antibody-to-DBCO ratio was measured via UV-Vis (Nanodrop)^28^. 1-2 µg DBCO-conjugated antibodies were combined with an equal volume of 80 µM azide-functionalized PLA oligomer (IDT) dissolved in PBS (Life Technologies) and allowed to react overnight at 4°C. The reaction mixture was then brought to 50% glycerol/PBS (Sigma Aldrich) for long-term storage at -20°C.

### Cell culture

Jurkat and Raji cell lines were a generous gift from Jun Huang^29^. Both were maintained at 37°C with 5% CO_2_ in RPMI (Gibco, Thermo Scientific) supplemented with 10% Fetal Bovine Serum (FBS, Hyclone, Fischer Scientific).

### Flow cytometry

Jurkat and Raji cells were plated in a 96-well plate (Corning) at 100,000 cells/well. Cells were centrifuged at 500g for 5 min, media was removed and replaced with 30 µL 5 nM PLA probes (2.5 nM probe A and 2.5 nM probe B) in Probe Binding Buffer (PBS, 0.1% BSA (Thermo Scientific), 0.1 mg/ml sonicated salmon sperm DNA (Invitrogen), 6.7 nM of each isotype control (mIgG1, mIgG2a, mIgG2b, and goat polyclonal)). Cell were incubated with probe for 30 min at 37°C. Cells were then washed three times by centrifuging at 500g for 5 min and resuspending in 100 µL 1% BSA/PBS. Cells were then resuspended in a 1 to 100 dilution of secondary antibody (Supplementary table 2) in 1% BSA/PBS and incubated for 20 min at room temperature. Cells were centrifuged and washed two times as before. Finally, they were analyzed using a Fortessa 4-15 (BD Biosciences) with a High Throughput Screening module.

### Drop-seq procedure

150,000 Jurkat and 150,000 Raji cells were counted and spun at 500 g for 3 min, washed once with 1% BSA/PBS, and spun again. Cells were then combined and plated in 96-well U-bottom plates. Cells were then centrifuged at 300 g for 3 min and fixed with 4 mM 3,3’– dithiobis(sulfosuccinimidyl propionate) (DTSSP, Life Technologies) in PBS at 37°C for 30 min. Cells were washed once with 1% BSA/PBS and 90µL probes were added at 5 nM each (2.5 nM probe A + 2.5 nM probe B) in Probe Binding Buffer. Cells were then incubated at 37°C for 60 min, washed twice with 1% BSA/PBS, and ligated with 300 µL Ligation Solution. Finally, cells were unfixed with 30 mM DTT at 37°C for 30 min, centrifuged at 300 g for 3 min, and resuspended in 0.1% BSA/PBS for Drop-seq separation.

### Smart-seq2 procedure

Cells underwent different processing procedures depending on whether they were analyzed by via a Drop-seq protocol or Smart-seq2 protocol. For Smart-seq2, 50,000 Jurkat cells and 50,000 Raji cells were collected, centrifuged at 500 g for 3 min, and washed once in 1% BSA/PBS. Jurkat cells were resuspended in 5 µM carboxyfluorescein diacetate succinimidyl ester (CFSE, Biolegend) in PBS for 20 min at room temperature in order to identify Jurkats specifically during cell sorting. The cells were then resuspended in 30 µL PLA probes in Probe Binding Buffer. Each probe was at 5 nM (2.5 nM probe A + 2.5 nM probe B). Cells were incubated at 37°C for 60 min. They were then centrifuged and washed three times by centrifuging and resuspending in 1% BSA/PBS at before. Cells were then resupended in 100 µL Ligase Solution (50mM HEPES pH 7.5, 10 mM MgCl_2_, 1mM rATP (New England Biolabs), 9.5 nM Connector oligomer (TTTCACGACACGACACGATTTAGGTC) (IDT), 130 U/ml T4 Ligase (NEB)) and rotated for 3 hr at 37°C. With 30 min remaining in the incubation, propidium iodide (Invitrogen) was added to the solution to a final concentration of 1 µg/ml. Cell were then centrifuged at 500g for 3 min, resuspended in 1% BSA/PBS, and sorted.

After PLA, cells were processed according to Drop-seq or Smart-seq2 protocol with some modifications. Details are available in the Supplementary Methods.

### Next-generation sequencing

For the Drop-seq pipeline, a NextSeq Mid-output kit v2 was used to sequence both mRNA and PLA libraries in the same sequencing run. The cDNA and PLA libraries each received 20% of the total reads. PhiX control was spiked in at 40% concentration according to Illumina’s instruction, because of the low diversity of the PLA libraries. Custom read 1 sequencing primer (Read1CustomSeqB), custom read 2 primer (DropPLA_Read2) and custom i7 index read primer (DropPLA_i7Read) were used according to Illumina’s instruction. Read distribution was 20 bases for read 1, 85 bases for read 2, and 8 bases for i7 index read.

For the Smart-seq2 pipeline, only the PLA library was sequenced. A mid-output NextSeq kit v2 was sufficient to sequence the pooled PLA libraries from four 96-well plates. PhiX control was spiked in at 40% concentration according to Illumina’s instruction. Custom read 1 sequencing primer (SmartPLA_Read1), custom i5 index read primer (SmartPLA_i5Read) and custom i7 index read primer (SmartPLA_i7Read) were used according to Illumina’s instruction. Read distribution was 75 bases for read 1, 8 bases for i5 index read, and 8 bases for i7 index read.

### Sequencing alignment

Sequencing data was aligned using a Java program. Details are available in the Supplementary Methods.

