## Supplementary information for "Quantification of proteins, protein complexes and mRNA in single cells by proximity-sequencing"

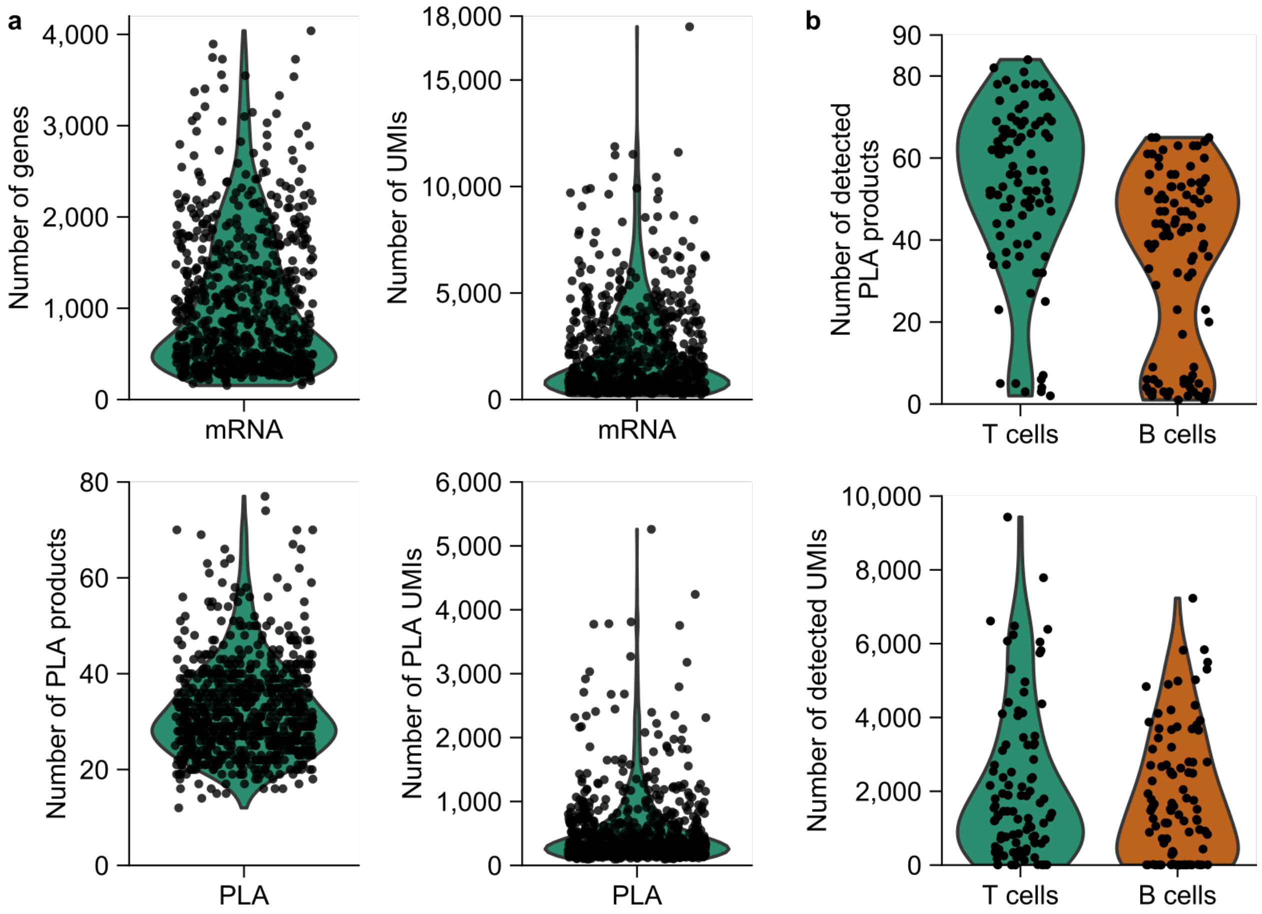


**Supplementary Fig. 1:** Quality analysis of Prox-seq data. **a,** Violin plots showing the number of detected genes and PLA products, and number of detected mRNA and PLA UMIs from Drop-seq pipeline. **b,** Violin plots showing the number of detected PLA products and PLA UMIs from Smart-seq2 pipeline.

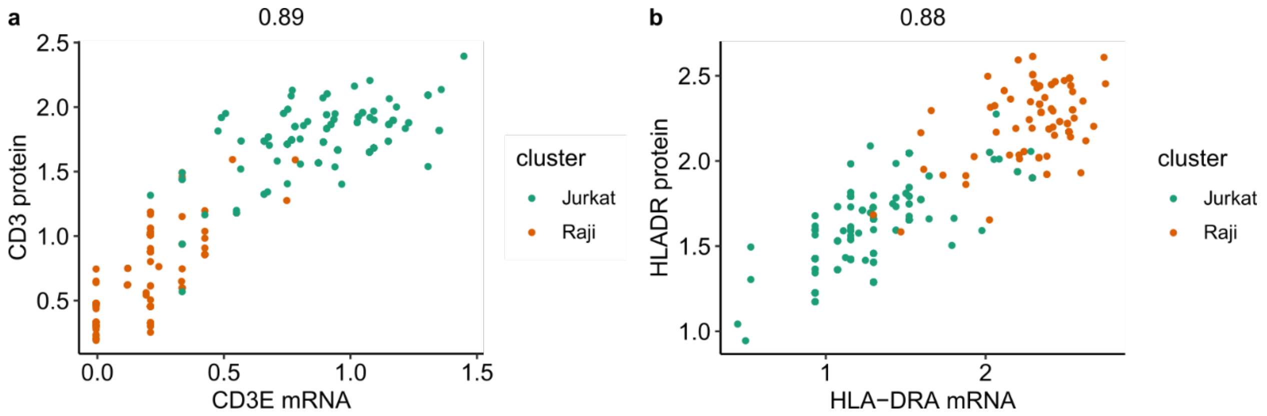


**Supplementary Fig. 2:** Correlation between protein and mRNA data measured by Prox-seq. **a**, Aggregated normalized expression levels of CD3E gene and CD3 protein. **b**, Aggregated normalized expression levels of HLA-DRA gene and HLA-DR protein. Each point represents the normalized expression level of the gene/protein averaged across each single cell's 5 nearest neighbors. The clusters were identified from the mRNA data. The numbers above the figures indicate the Pearson correlation coefficient.

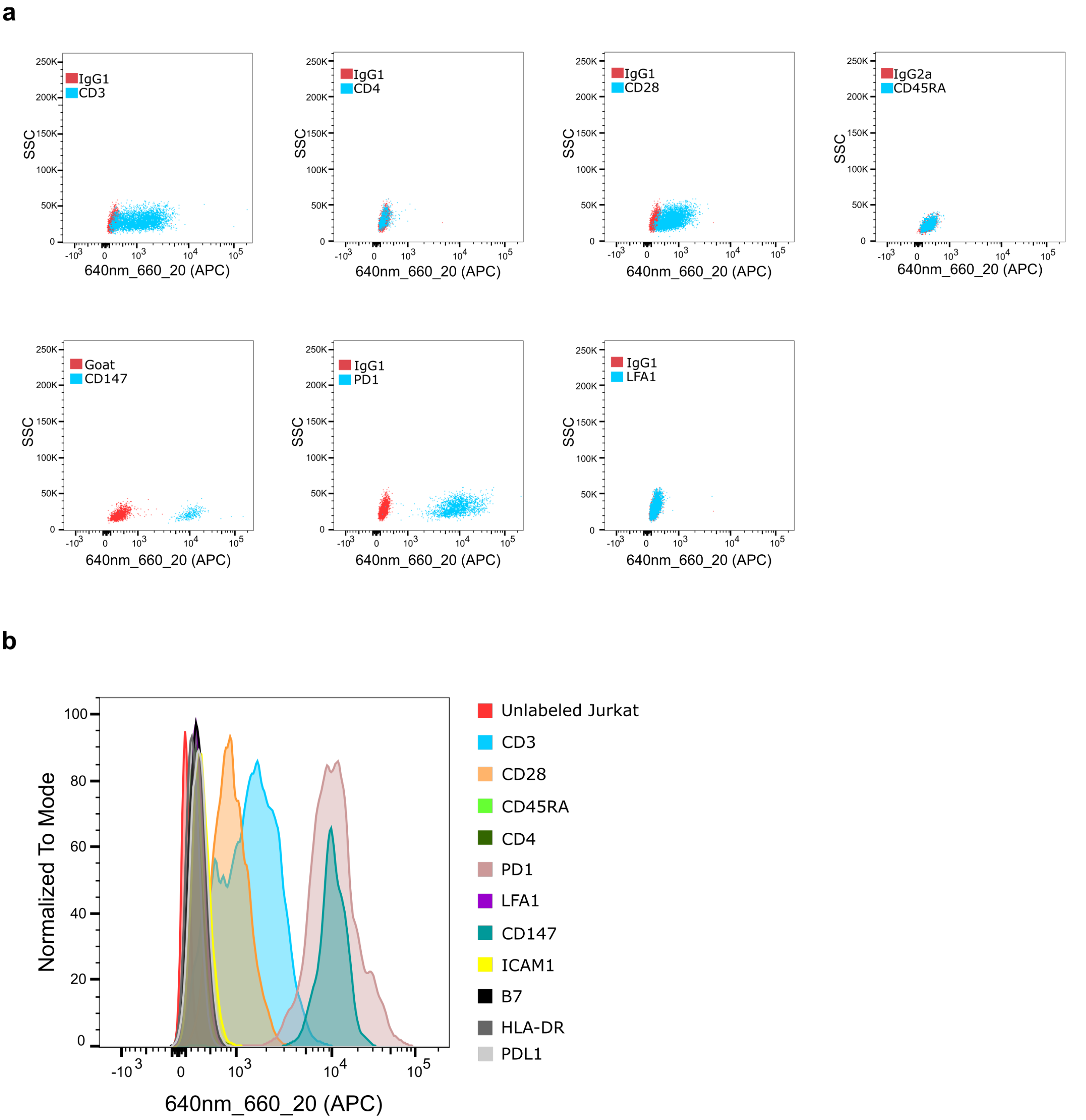


**Supplementary Fig. 3:** Flow cytometry data showing PLA probe binding on Jurkat cells. **a,** Each T-cell marker in the panel compared to its corresponding isotype control. **b,** The amount of binding for all PLA probes on Jurkat cells.

**a**

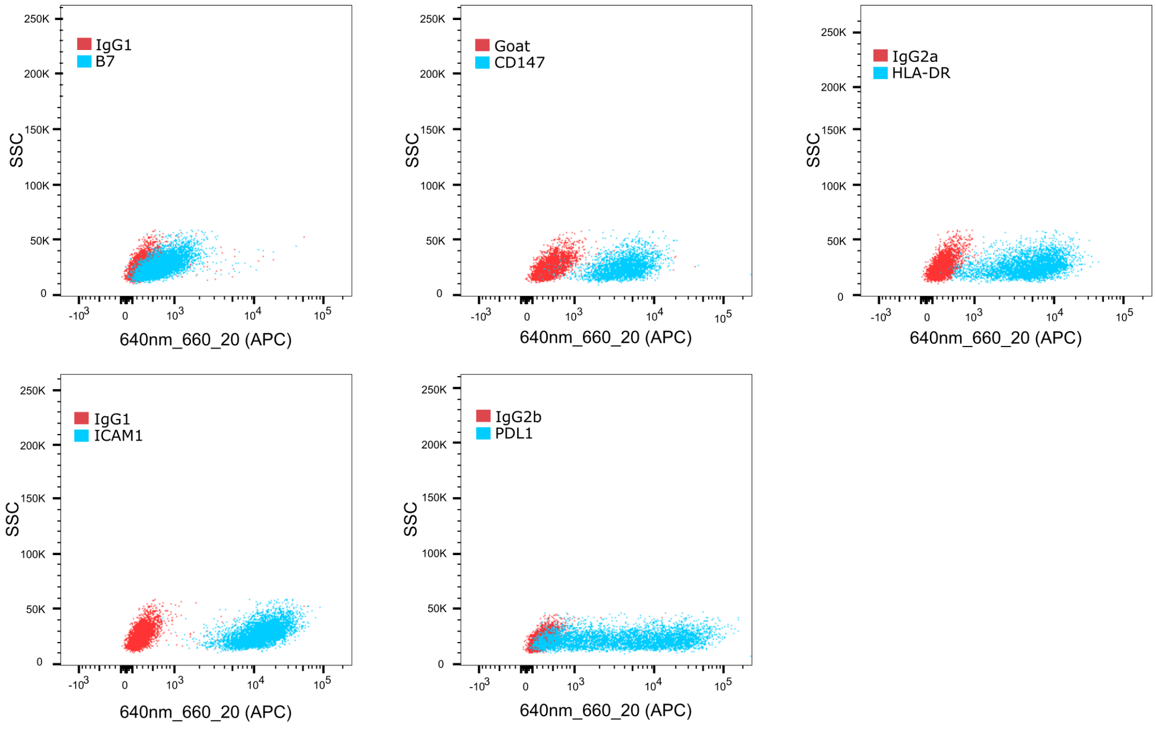


**b**

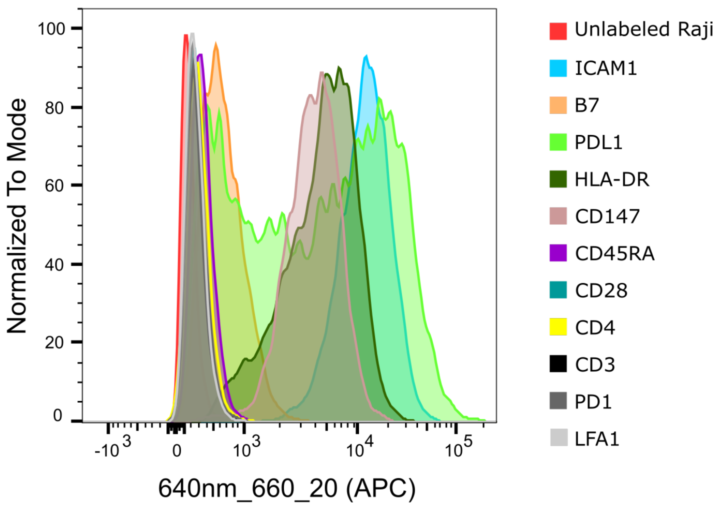


**Supplementary Fig. 4:** Flow cytometry data showing PLA probe binding on Raji cells. **a,** Each B-cell marker in the panel compared to its corresponding isotype control. **b,** The amount of binding for all PLA probes on Raji cells.

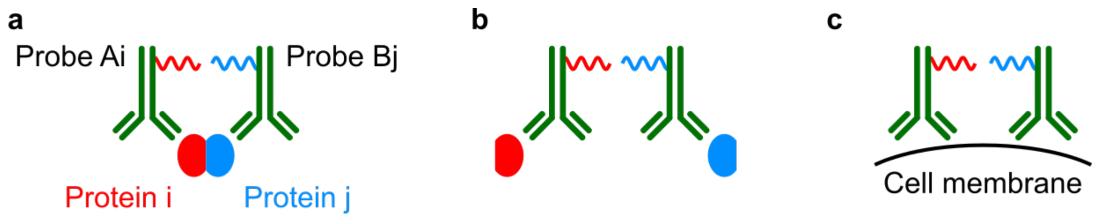


**Supplementary Fig. 5:** Three possible sources of PLA products. (a) Ligation of probes targeting the same protein complex. (b) Ligation of probes that are in proximity by chance. (c) Ligation of probes that bind non-specifically, and in close proximity to each other.

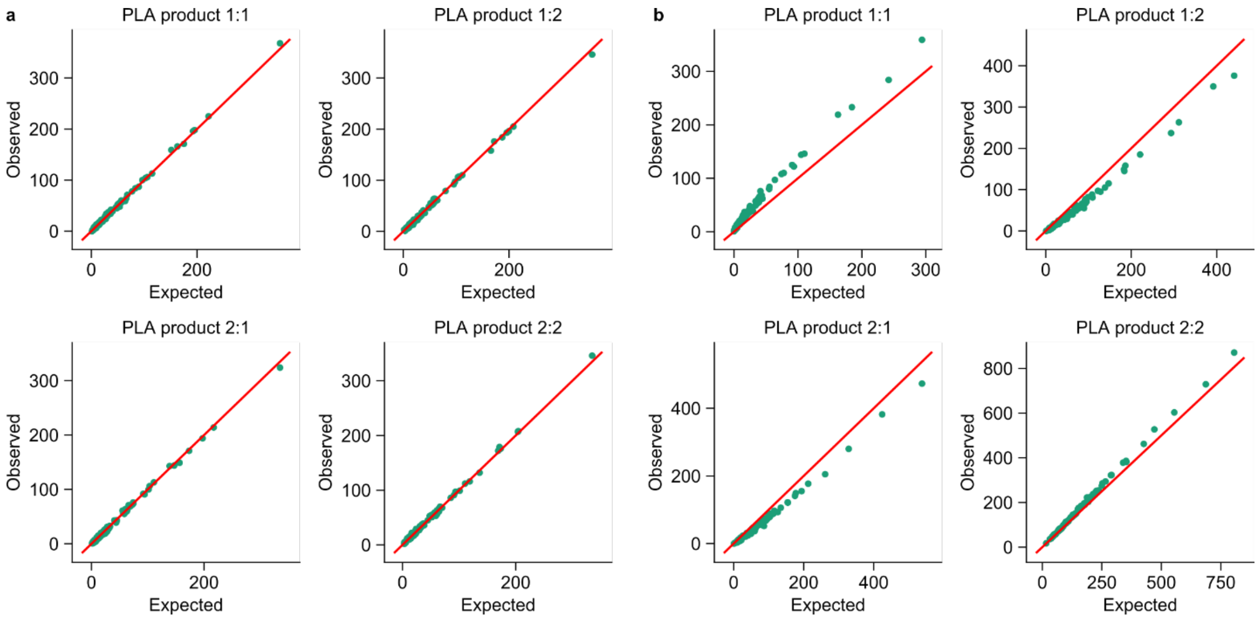


**Supplementary Fig. 6:** Example simulated PLA data from 2 protein targets. **a-b,** Scatter plots showing the observed PLA count against the expected PLA count when **(a)** no protein complex exists, and when **(b)** 2:2 is a homodimer**.** Each dot represents a single cell (n=100 cells). The red line indicates the y=x line.

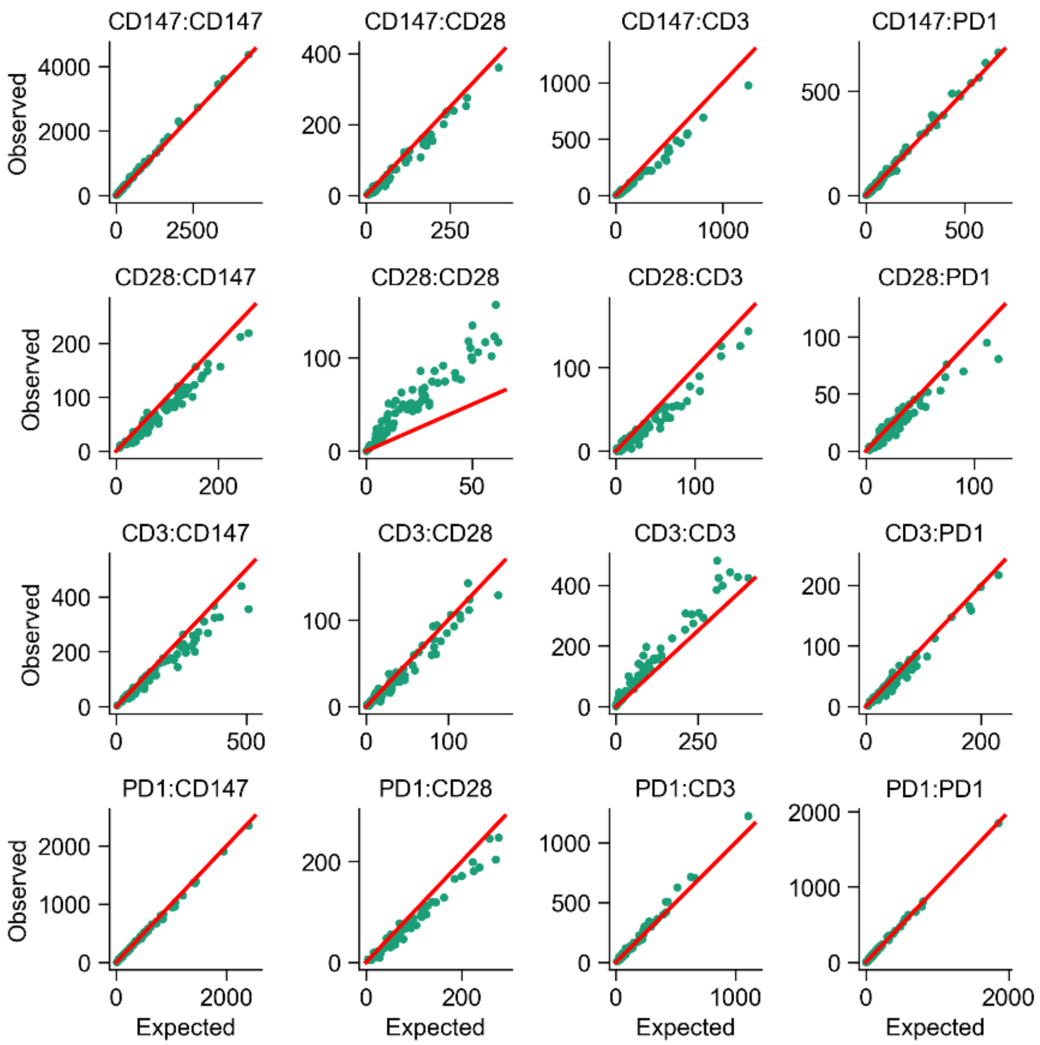


**Supplementary Fig. 7:** Plot of expected and observed PLA count of T cell markers among T cells. The red line indicates the y=x line.

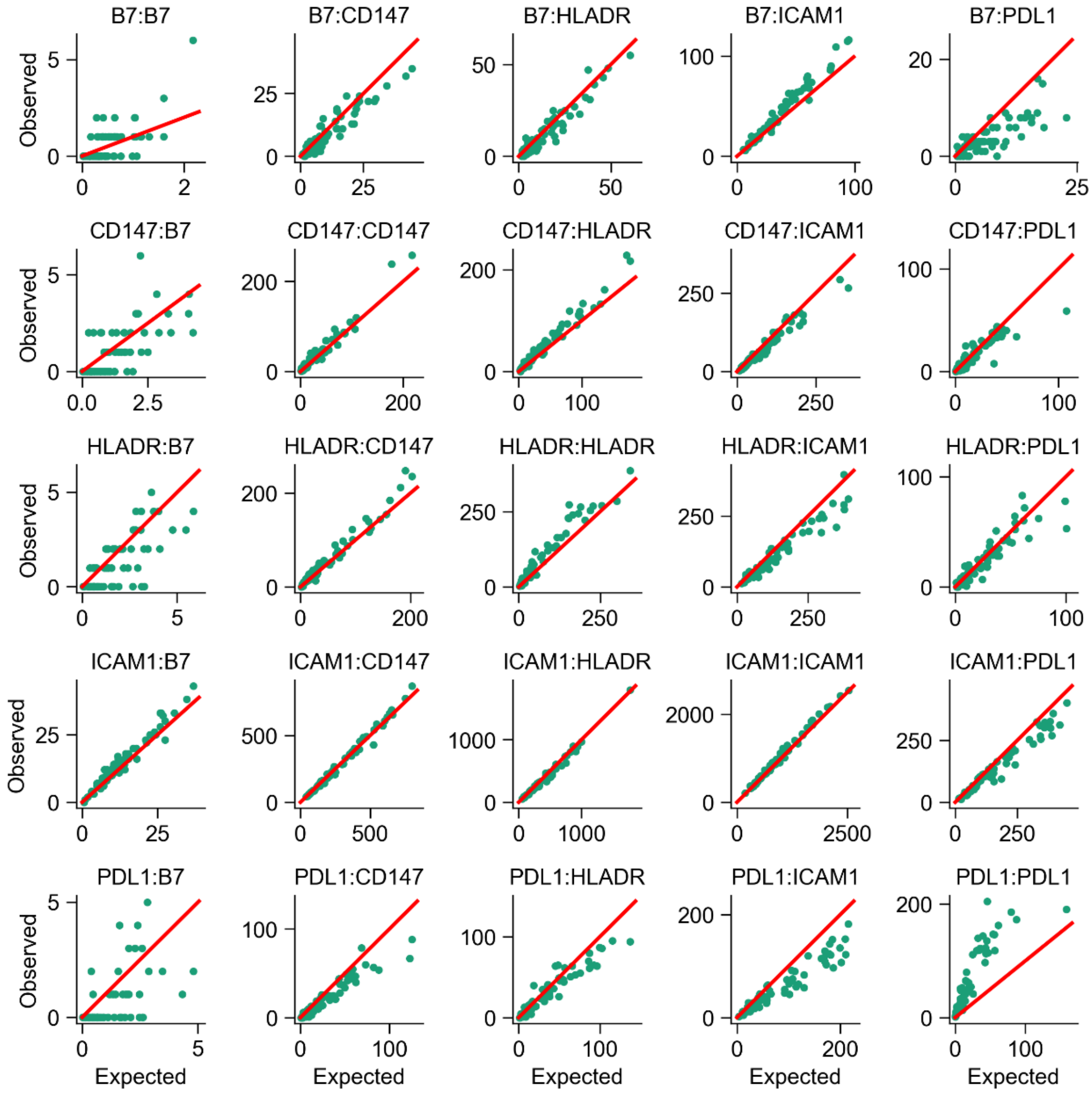


**Supplementary Fig. 8:** Plot of expected and observed PLA count of B cell markers among B cells. The red line indicates the y=x line.

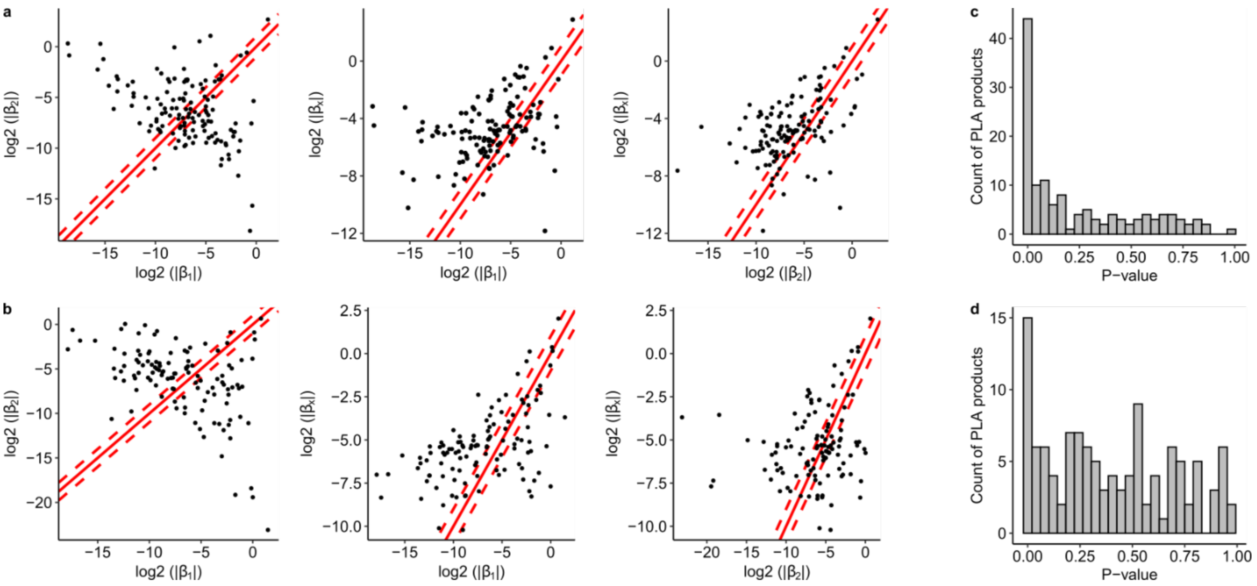


**Supplementary Fig. 9:** Linear model of PLA product formation. **a-b,** Scatter plots of the magnitude of the effect of probe abundance (β1 and β2) and the interactions term (βx) for all PLA products detected from **(a)** T cells and **(b)** B cells. **c-d,** Histograms of the P-values of the likelihood ratio test for **(c)** T cells and **(d)** B cells. In **(a, b)**, the red solid lines indicate equal magnitude, and the red dashed lines indicate a two-fold difference in magnitude.

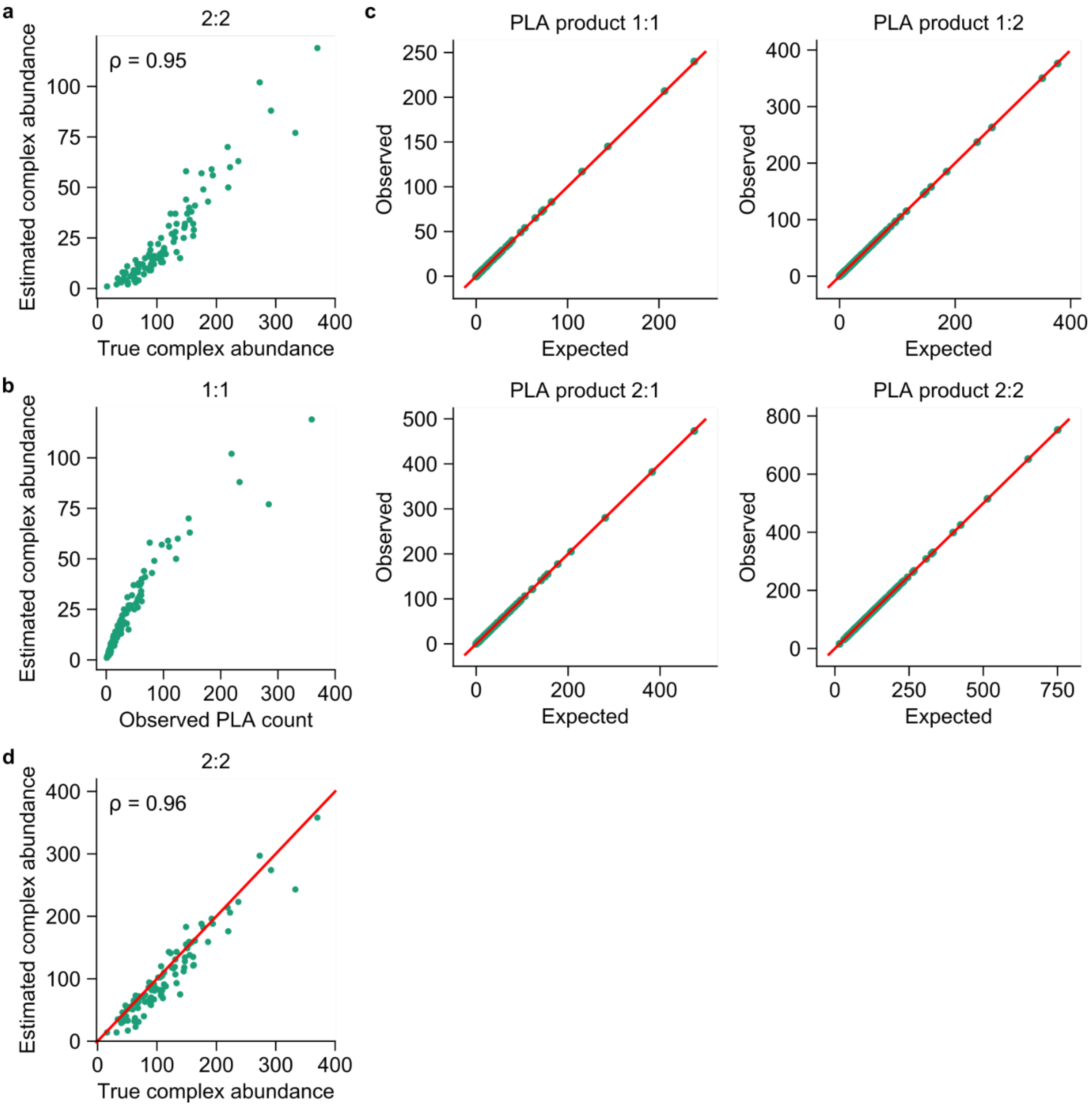


**Supplementary Fig. 10:** Estimated complex abundance in simulated data. (a) Plot of the estimated abundance of the true positive complex 2:2 against its true complex abundance from the simulated data shown in Fig. 2b. (b) Plot of the estimated abundance of the false positive complex 1:1 against its observed PLA count from the same simulated data set. (c) Scatter plots showing the observed count against the expected count, after adjusting for the complex abundance. (d) Plot of the estimated abundance of complex 2:2 against its true complex abundance, when 1:1 is forced to not be a complex. Each dot represents a single cell (n=100 cells). The red lines in (c) and (d) indicate the y=x line.

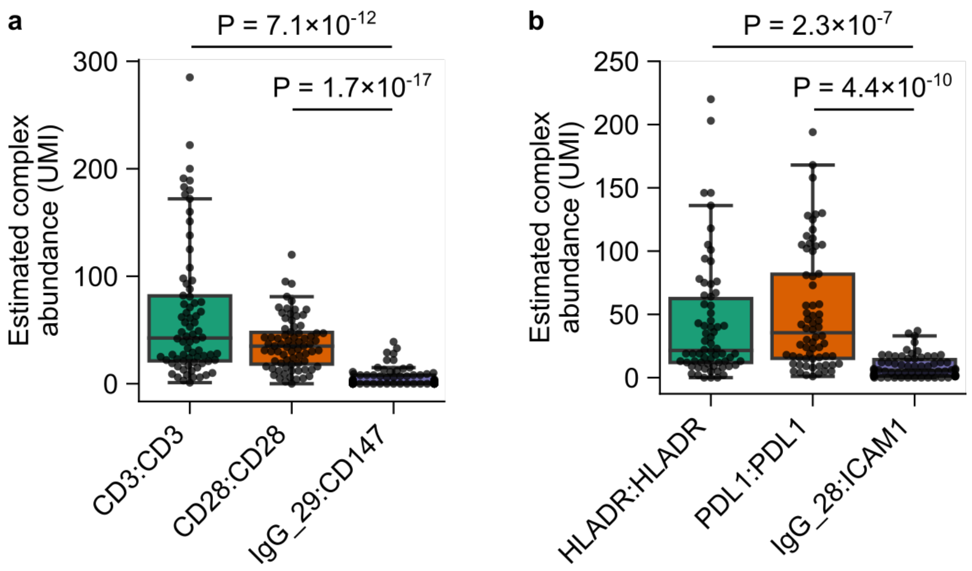


**Supplementary Fig. 11:** Estimated complex abundance compared to negative control PLA products. **a,** Estimated abundance of CD3 and CD28 homodimers compared to an isotype control complex in Jurkat cells. **b,** Estimated abundance of HLA-DR and PDL1 homodimers compared to an isotype control complex in Raji cells cells. The box shows the quartiles of the data, and the whiskers extend by 1.5× the interquartile range beyond the box. P values are calculated using two-tailed Welch's t-test, without multiple comparison correction.

| Target | Clone | Species | Company | Catalog Number | Barcode Drop-seq | Barcode Smart-seq2 |
|---|---|---|---|---|---|---|
| CD3 | UCHT1 | Mouse | Biolegend | 300437 | ATAGCGTC | ATAGCGTC |
| CD28 | CD28.2 | Mouse | Biolegend | 302933 | CAAGGTCT | CAAGGTCT |
| CD45RA | HI100 | Mouse | Biolegend | 304102 | CAATCGGT | CCACAATG |
| CD4 | OKT4 | Mouse | Biolegend | 317402 | GCATGTAC | GCATGTAC |
| PD1 | NAT105 | Mouse | Biolegend | 367402 | TAAGGCTC | TAAGGCTC |
| LFA1 | m24 | Mouse | Biolegend | 363402 | CTAGCTGA | CTAGCTGA |
| ICAM1 | HA58 | Mouse | Biolegend | 353102 | ATACGTGC | ATACGTGC |
| B7 | 2D10 | Mouse | Biolegend | 305202 | AGACCTTG | AGACCTTG |
| PDL1 | 29E.2A3 | Mouse | Biolegend | 329702 | GTAGCACT | GTAGCACT |
| HLA-DR | L243 | Mouse | Biolegend | 307602 | AGAAGAGG | AGAAGAGG |
| CD147 | Polyclonal | Goat | R&D Systems | AF972 | TTACGGTG | GCAGTATC |
| Isotype control | Polyclonal | Mouse | EMD Millipore | PP54 | | TGAGACCT |
| Isotype control | Polyclonal | Goat | R&D Systems | AB-108-C | | GAAGGAAG |
| Isotype control | MG1-45 | Mouse IgG1 | Biolegend | 401401 | CGATTGAC | |
| Isotype control | MG2a-53 | Mouse IgG2a | Biolegend | 401501 | ACAGTGCT | |
| Isotype control | 27-35 | Mouse IgG2b | Biolegend | 402211 | TAACTGGC | |

**Supplementary Table 1. List of all antibodies used to make PLA probes**

| Target | Species | Company | Catalog number | Fluorphore |
|---|---|---|---|---|
| Mouse | Donkey | R&D Systems | NL008 | NL637 |
| Goat | Donkey | R&D Systems | NL002 | NL637 |

**Supplementary Table 2. List of secondary antibodies**

| Name | Sequence | Note |
|---|---|---|
| TSO_RNAhybrid | AAGCAGTGGTATCAACGCAGAGTGAA TrGrGrG | For template switching |
| TSO_PCR | AAGCAGTGGTATCAACGCAGAGT | For amplifying cDNAs |
| U_fwd | GCATCGTCTCGTGGGCTC | For amplifying PLA products |
| P5_TSO | AATGATACGGCGACCACCGAGATCTA CACGCCT GTCCGCGGAAGCAGTGGTATCAACGC AGAGT* A*C | For adding P5 adapter |
| P7_N7XX_Custom 2 | CAAGCAGAAGACGGCATACGAGAT**[i7]** AGATGCATCGTCTCGTGGGCTCGG<br>i7= TCGCCTTA for N701<br>i7= CTAGTACG for N702<br>i7= TTCTGCCT for N703<br>i7= GCTCAGGA for N704 | For adding P7 adapter and i7 index |
| Read1CustomSeq B | GCCTGTCCGCGGAAGCAGTGGTATCA ACGCAGAGTAC | Custom read 1 sequencing primer |
| DropPLA_Read2 | AGATGCATCGTCTCGTGGGCTCGG | Custom read 2 sequencing primer |
| DropPLA_i7Read | CCGAGCCCACGAGACGATGCATCT | Custom i7 index sequencing primer |

**Supplementary Table 3: List of primers used in the Drop-seq protocol**

| Name | Sequence | Note |
|---|---|---|
| Oligo_dTGT | AAGCAGTGGTATCAACGCAGAGTACT TTTTTTTTTTTTTTTTTTTTTTTTTTTTTG T | For priming the poly-A tail of PLA products |
| U_fwd | GCATCGTCTCGTGGGCTC | For amplifying PLA products (same as U_fwd used in Drop-seq pipeline) |
| SmartPLA_P5_S5XX | AATGATACGGCGACCACCGAGATCTACAC**[i5]**TATTTGGCATCGTCTCGTGGGCTCGG<br>List of full sequences are in Supplementary Table 5 | For adding P5 adapter and i5 index |
| SmartPLA_P7_N7XX | CAAGCAGAAGACGGCATACGAGAT**[i7]**GCCGAAGCAGTGGTATCAACGCAGAGT<br>List of full sequences are in Supplementary Table 5 | For adding P7 adapter and i7 index |
| SmartPLA_Read1 | TATTTGGCATCGTCTCGTGGGCTCGG | Custom read 1 sequencing primer |
| SmartPLA_i5Read | CCGAGCCCACGAGACGATGCCAAATA | Custom i5 index sequencing primer |
| SmartPLA_i7Read | ACTCTGCGTTGATACCACTGCTTCGGC | Custom i7 index sequencing primer |

**Supplementary Table 4: List of primers used in the Smart-seq2 protocol**

| Primer sequence | Primer sequence | Index number | Index sequence |
|---|---|---|---|
| SmartPLA_ P7_N701 | CAAGCAGAAGACGGCATACGAGATTCGCCT TAGCCGAAGCAGTGGTATCAACGCAGAGT | N701 | TCGCCTTA |
| SmartPLA_ P7_N702 | CAAGCAGAAGACGGCATACGAGATCTAGTA CGGCCGAAGCAGTGGTATCAACGCAGAGT | N702 | CTAGTACG |
| SmartPLA_ P7_N703 | CAAGCAGAAGACGGCATACGAGATTTCTGC CTGCCGAAGCAGTGGTATCAACGCAGAGT | N703 | TTCTGCCT |
| SmartPLA_ P7_N704 | CAAGCAGAAGACGGCATACGAGATGCTCAG GAGCCGAAGCAGTGGTATCAACGCAGAGT | N704 | GCTCAGGA |
| SmartPLA_ P7_N706 | CAAGCAGAAGACGGCATACGAGATCATGCC TAGCCGAAGCAGTGGTATCAACGCAGAGT | N706 | CATGCCTA |
| SmartPLA_ P7_N707 | CAAGCAGAAGACGGCATACGAGATGTAGAG AGGCCGAAGCAGTGGTATCAACGCAGAGT | N707 | GTAGAGAG |
| SmartPLA_ P7_N708 | CAAGCAGAAGACGGCATACGAGATCCTCTCT GGCCGAAGCAGTGGTATCAACGCAGAGT | N708 | CCTCTCTG |
| SmartPLA_ P7_N709 | CAAGCAGAAGACGGCATACGAGATAGCGTA GCGCCGAAGCAGTGGTATCAACGCAGAGT | N709 | AGCGTAGC |
| SmartPLA_ P7_N710 | CAAGCAGAAGACGGCATACGAGATCAGCCT CGGCCGAAGCAGTGGTATCAACGCAGAGT | N710 | CAGCCTCG |
| SmartPLA_ P7_N711 | CAAGCAGAAGACGGCATACGAGATTGCCTC TTGCCGAAGCAGTGGTATCAACGCAGAGT | N711 | TGCCTCTT |
| SmartPLA_ P7_N712 | CAAGCAGAAGACGGCATACGAGATTCCTCTA CGCCGAAGCAGTGGTATCAACGCAGAGT | N712 | TCCTCTAC |
| SmartPLA_ P7_N715 | CAAGCAGAAGACGGCATACGAGATCCTGAG ATGCCGAAGCAGTGGTATCAACGCAGAGT | N715 | CCTGAGAT |
| SmartPLA_ P7_N716 | CAAGCAGAAGACGGCATACGAGATTAGCGA GTGCCGAAGCAGTGGTATCAACGCAGAGT | N716 | TAGCGAGT |
| SmartPLA_ P7_N718 | CAAGCAGAAGACGGCATACGAGATGTAGCT CCGCCGAAGCAGTGGTATCAACGCAGAGT | N718 | GTAGCTCC |
| SmartPLA_ P7_N719 | CAAGCAGAAGACGGCATACGAGATTACTAC GCGCCGAAGCAGTGGTATCAACGCAGAGT | N719 | TACTACGC |
| SmartPLA_ P7_N720 | CAAGCAGAAGACGGCATACGAGATAGGCTC CGGCCGAAGCAGTGGTATCAACGCAGAGT | N720 | AGGCTCCG |
| SmartPLA_ P7_N721 | CAAGCAGAAGACGGCATACGAGATGCAGCG TAGCCGAAGCAGTGGTATCAACGCAGAGT | N721 | GCAGCGTA |
| SmartPLA_ P7_N722 | CAAGCAGAAGACGGCATACGAGATCTGCGC ATGCCGAAGCAGTGGTATCAACGCAGAGT | N722 | CTGCGCAT |
| SmartPLA_ P7_N723 | CAAGCAGAAGACGGCATACGAGATGAGCGC TAGCCGAAGCAGTGGTATCAACGCAGAGT | N723 | GAGCGCTA |
| SmartPLA_ P7_N724 | CAAGCAGAAGACGGCATACGAGATCGCTCA GTGCCGAAGCAGTGGTATCAACGCAGAGT | N724 | CGCTCAGT |
| SmartPLA_ P7_N726 | CAAGCAGAAGACGGCATACGAGATGTCTTA GGGCCGAAGCAGTGGTATCAACGCAGAGT | N726 | GTCTTAGG |

| | | | |
|---|---|---|---|
| SmartPLA_ P7_N727 | CAAGCAGAAGACGGCATACGAGATACTGAT CGGCCGAAGCAGTGGTATCAACGCAGAGT | N727 | ACTGATCG |
| SmartPLA_ P7_N728 | CAAGCAGAAGACGGCATACGAGATTAGCTG CAGCCGAAGCAGTGGTATCAACGCAGAGT | N728 | TAGCTGCA |
| SmartPLA_ P7_N729 | CAAGCAGAAGACGGCATACGAGATGACGTC GAGCCGAAGCAGTGGTATCAACGCAGAGT | N729 | GACGTCGA |

**Supplementary Table 5: List of indexing primers used in Smart-seq2 protocol**

| PLA Oligomer Name | Sequence | Barcode |
|---|---|---|
| PLA.Seq.Bead.9.A | /5AzideN/CGCATTGCATCGTCTCGTGGGCTCGGCHHHHACHHHHACHHHNGCAGATAGCGTCGATCGCTAAATCGTG | ATAGCGTC |
| PLA.Seq.Bead.9.B | /5Phos/TCGTGTCGTGTCTAAAGTCCATAGCGTCACAAAAAAAAAAAAAAAAAAAAAAAAAAAAA/3AzideN/ | ATAGCGTC |
| PLA.Seq.Bead.10.A | /5AzideN/CGCATTGCATCGTCTCGTGGGCTCGGCHHHHACHHHHACHHHNGCAGAGACCTTGGATCGCTAAATCGTG | AGACCTTG |
| PLA.Seq.Bead.10.B | /5Phos/TCGTGTCGTGTCTAAAGTCCAGACCTTGACAAAAAAAAAAAAAAAAAAAAAAAAAAAAA/3AzideN/ | AGACCTTG |
| PLA.Seq.Bead.11.A | /5AzideN/CGCATTGCATCGTCTCGTGGGCTCGGCHHHHACHHHHACHHHNGCAGCCACAATGGATCGCTAAATCGTG | CCACAATG |
| PLA.Seq.Bead.11.B | /5Phos/TCGTGTCGTGTCTAAAGTCCCCACAATGACAAAAAAAAAAAAAAAAAAAAAAAAAAAAA/3AzideN/ | CCACAATG |
| PLA.Seq.Bead.12.A | /5AzideN/CGCATTGCATCGTCTCGTGGGCTCGGCHHHHACHHHHACHHHNGCAGTGAGACCTGATCGCTAAATCGTG | TGAGACCT |
| PLA.Seq.Bead.12.B | /5Phos/TCGTGTCGTGTCTAAAGTCCTGAGACCTACAAAAAAAAAAAAAAAAAAAAAAAAAAAAA/3AzideN/ | TGAGACCT |
| PLA.Seq.Bead.13.A | /5AzideN/CGCATTGCATCGTCTCGTGGGCTCGGCHHHHACHHHHACHHHNGCAGGTAGCACTGATCGCTAAATCGTG | GTAGCACT |
| PLA.Seq.Bead.13.B | /5Phos/TCGTGTCGTGTCTAAAGTCCGTAGCACTACAAAAAAAAAAAAAAAAAAAAAAAAAAAAA/3AzideN/ | GTAGCACT |
| PLA.Seq.Bead.14.A | /5AzideN/CGCATTGCATCGTCTCGTGGGCTCGGCHHHHACHHHHACHHHNGCAGCAATCGGTGATCGCTAAATCGTG | CAATCGGT |
| PLA.Seq.Bead.14.B | /5Phos/TCGTGTCGTGTCTAAAGTCCCAATCGGTACAAAAAAAAAAAAAAAAAAAAAAAAAAAAA/3AzideN/ | CAATCGGT |
| PLA.Seq.Bead.15.A | /5AzideN/CGCATTGCATCGTCTCGTGGGCTCGGCHHHHACHHHHACHHHNGCAGCTAGCTGAGATCGCTAAATCGTG | CTAGCTGA |
| PLA.Seq.Bead.15.B | /5Phos/TCGTGTCGTGTCTAAAGTCCCTAGCTGAACAAAAAAAAAAAAAAAAAAAAAAAAAAAAA/3AzideN/ | CTAGCTGA |
| PLA.Seq.Bead.17.A | /5AzideN/CGCATTGCATCGTCTCGTGGGCTCGGCHHHHACHHHHACHHHNGCAGGAAGGAAGGATCGCTAAATCGTG | GAAGGAAG |
| PLA.Seq.Bead.17.B | /5Phos/TCGTGTCGTGTCTAAAGTCCGAAGGAAGACAAAAAAAAAAAAAAAAAAAAAAAAAAAAA/3AzideN/ | GAAGGAAG |
| PLA.Seq.Bead.18.A | /5AzideN/CGCATTGCATCGTCTCGTGGGCTCGGCHHHHACHHHHACHHHNGCAGAGAAGAGGGATCGCTAAATCGTG | AGAAGAGG |
| PLA.Seq.Bead.18.B | /5Phos/TCGTGTCGTGTCTAAAGTCCAGAAGAGGACAAAAAAAAAAAAAAAAAAAAAAAAAAAAA/3AzideN/ | AGAAGAGG |
| PLA.Seq.Bead.19.A | /5AzideN/CGCATTGCATCGTCTCGTGGGCTCGGCHHHHACHHHHACHHHNGCAGTAAGGCTCGATCGCTAAATCGTG | TAAGGCTC |

| | | |
|---|---|---|
| PLA.Seq.Bead.19.B | /5Phos/TCGTGTCGTGTCTAAAGTCCTAAGGCTCACAAAAAAAAAAAAAAAAAAAAAAAAAAAAA/3AzideN/ | TAAGGCTC |
| PLA.Seq.Bead.20.A | /5AzideN/CGCATTGCATCGTCTCGTGGGCTCGGCHHHHACHHHHACHHHNGCAGTTACGGTGGATCGCTAAATCGTG | TTACGGTG |
| PLA.Seq.Bead.20.B | /5Phos/TCGTGTCGTGTCTAAAGTCCTTACGGTGACAAAAAAAAAAAAAAAAAAAAAAAAAAAAA/3AzideN/ | TTACGGTG |
| PLA.Seq.Bead.21.A | /5AzideN/CGCATTGCATCGTCTCGTGGGCTCGGCHHHHACHHHHACHHHNGCAGGCATGTACGATCGCTAAATCGTG | GCATGTAC |
| PLA.Seq.Bead.21.B | /5Phos/TCGTGTCGTGTCTAAAGTCCGCATGTACACAAAAAAAAAAAAAAAAAAAAAAAAAAAAA/3AzideN/ | GCATGTAC |
| PLA.Seq.Bead.22.A | /5AzideN/CGCATTGCATCGTCTCGTGGGCTCGGCHHHHACHHHHACHHHNGCAGCAAGGTCTGATCGCTAAATCGTG | CAAGGTCT |
| PLA.Seq.Bead.22.B | /5Phos/TCGTGTCGTGTCTAAAGTCCCAAGGTCTACAAAAAAAAAAAAAAAAAAAAAAAAAAAAA/3AzideN/ | CAAGGTCT |
| PLA.Seq.Bead.23.A | /5AzideN/CGCATTGCATCGTCTCGTGGGCTCGGCHHHHACHHHHACHHHNGCAGATACGTGCGATCGCTAAATCGTG | ATACGTGC |
| PLA.Seq.Bead.23.B | /5Phos/TCGTGTCGTGTCTAAAGTCCATACGTGCACAAAAAAAAAAAAAAAAAAAAAAAAAAAAA/3AzideN/ | ATACGTGC |
| PLA.Seq.Bead.28.A | /5AzideN/CGCATTGCATCGTCTCGTGGGCTCGGCHHHHACHHHHACHHHNGCAGCGATTGACGATCGCTAAATCGTG | CGATTGAC |
| PLA.Seq.Bead.28.B | /5Phos/TCGTGTCGTGTCTAAAGTCCCGATTGACACAAAAAAAAAAAAAAAAAAAAAAAAAAAAA/3AzideN/ | CGATTGAC |
| PLA.Seq.Bead.29.A | /5AzideN/CGCATTGCATCGTCTCGTGGGCTCGGCHHHHACHHHHACHHHNGCAGACAGTGCTGATCGCTAAATCGTG | ACAGTGCT |
| PLA.Seq.Bead.29.B | /5Phos/TCGTGTCGTGTCTAAAGTCCACAGTGCTACAAAAAAAAAAAAAAAAAAAAAAAAAAAAA/3AzideN/ | ACAGTGCT |
| PLA.Seq.Bead.30.A | /5AzideN/CGCATTGCATCGTCTCGTGGGCTCGGCHHHHACHHHHACHHHNGCAGTAACTGGCGATCGCTAAATCGTG | TAACTGGC |
| PLA.Seq.Bead.30.B | /5Phos/TCGTGTCGTGTCTAAAGTCCTAACTGGCACAAAAAAAAAAAAAAAAAAAAAAAAAAAAA/3AzideN/ | TAACTGGC |

**Supplementary Table 6: List of all oligomers used to make PLA probes**

### SUPPLEMENTARY METHODS

#### Drop-seq pipeline experiment

After the PLA process, cells were resuspended in 0.01% BSA at 100,000 cells/ml, injected into a 3 ml syringe and kept on ice until ready to use. Drop-seq beads (ChemGenes, cat # Macosko-2011-10(V+)) were washed once with TE-TW (10 mM Tris pH 8.0, 1 mM EDTA and 0.01% Tween-20), resuspended in Drop-seq lysis buffer at 120,000 beads/ml[1], injected into a 3 ml syringe along with a magnetic disc (V&P Scientific, cat # 772DP-N42-5-2) and kept on ice until ready to use. Droplet oil (Bio-Rad, cat # 1864006) was injected into a 10 ml syringe and kept at room temperature until ready to use.

The Drop-seq chip was made from PDMS using the design from Macosko *et. al.* 2015. The chip was mounted on an inverted microscope to monitor droplet formation. Then, the cell, bead and oil syringes were connected to the appropriate chip inlets via plastic tubes (Scientific Commodities Inc, cat # BB31695PE/2). A magnetic stirrer (V&P Scientific, cat # 710D2) was used to keep the beads resuspended during the experiment. To generate droplets, syringe pumps were used to inject cells at 3 ml/hour flow rate, beads at 3 ml/hour, and oil at 12 ml/hour. The generated droplets were collected in a 50 ml tube via a plastic tube that was inserted into the chip's outlet.

After droplet generation, the droplets were broken as follows. Oil at the bottom of the collection tube was removed as much as possible, without disturbing the droplets. Then, 30 ml of 6X Saline-Sodium Citrate solution (SSC, Fisher BioReagents) was added to the droplets, followed by 1ml of 1H,1H,2H,2H-Perfluorooctan-1-ol (Synquest Laboratories). Next, the tube was shaken strongly by hand 4 times, and centrifuged at 1000g for 1 minute. After this, the supernatant was transferred to another 50 ml tube (tube 1), and 30 ml of 6X SSC was added to the collection tube, and then the supernatant was transferred to another 50 ml tube (tube 2). Tubes 1 and 2 were centrifuged at 1000g for 3 minutes to pellet the beads, then the supernatant in both tubes was removed, and the beads in both tubes were transferred to a 1.5 ml microfuge tube. The beads were washed twice with 1 ml of 6X SSC, and once with 300 μL of 5X Maxima H- reverse transcription (RT) buffer (Thermo Scientific) before proceeding to RT.

The RT mixture was prepared as following: 1X RT buffer, 4% Ficoll PM-400 (Sigma-Aldrich), 1 mM each dNTP (NEB), 1,000 units/ml murine RNase inhibitor (NEB), 2.5 μM TSO_RNAhybrid (IDT), and 10 units/μL Maxima H- reverse transcriptase (Thermo Scientific). After bead washing, 200 μL of RT mixture was added to the beads, and the beads were incubated at room temperature for 30 minutes with rotation, followed by incubation at 42C for 90 minutes with rotation.

After RT, the beads were centrifuged at 1000g for 1 minute, the supernatant was discarded, and the beads were washed once with 1 ml of TE-SDS (10 mM Tris pH 8.0, 1 mM EDTA and 0.5% SDS), twice with 1 ml of TE-TW, and once with 1 ml of Tris pH 8.0. Then, 200 µL of exonuclease I mixture (1,000 units/ml exonuclease I and 1X exonuclease I buffer, NEB, cat # M0293) was added to the beads, and the beads were incubated at 37C for 45 minutes with rotation.

After exonuclease I treatment, the beads were centrifuged at 1000g for 1 minute, washed once with 1 ml of TE-SDS, twice with 1 ml of TE-TW, and once with 1 ml of nuclease-free water. Next, the cDNA and PLA products on the beads were amplified using PCR. The beads were distributed into a 96-well plate at approximately 5,000 beads per well. Then, a 50 µL PCR reaction was set up in each well, which contained 1X KAPA HiFi HotStart Readymix (Roche), 0.8 µM TSO_PCR primer, and 0.8 µM U_fwd primer. The thermal cycle program was: 95C 3 min; 4 cycles of 98C 20s, 65C 45s, 72C 3 min; 10 cycles of 98C 20s, 67C 20s, 72C 3 min; 72C 5 min; 4C hold.

After PCR, 10 µL of nuclease-free water was added to each well, the plate was centrifuged at 1000g for 1 minute, and 50 µL of supernatant from each well was transferred to another 96-well plate (plate 1). Then, 30 µL of AMPure XP beads (Beckman Coulter, cat # A63881) was added to each well (ie, 0.6X AMPure bead concentration), the new plate was incubated at room temperature for 5 minutes, and left on a magnetic rack for 5 minutes. Next, the supernatant, which contained PLA products, was transferred into another 96-well plate (plate 2). Each well in plate 1 was washed four times with 200 µL of 80% ethanol, and left to air dry for 2 minutes. 11.5 µL of nuclease-free water was added to each well to elute the cDNA products from the AMPure beads, the plate was incubated at room temperature for 5 minutes, left on a magnetic rack for 3 minutes, and 10 µL of the supernatant from each well was collected and pooled into a 1.5 ml microfuge tube. To recover the PLA products from the supernatant stored in plate 2, 60 µL of AMPure beads was added to each well (ie, 1.8X AMPure bead concentration), and the same protocol for plate 1 was followed afterwards.

cDNA products were quantified using High Sensitivity DNA TapeStation (Agilent). They have a broad length distribution between 400 to more than 3000 base pair, weak a peak at approximately 1000 bp. Library preparation of cDNA products was performed using Nextera XT DNA Library Preparation Kit (Illumina, cat # FC-131-1024). 450 pg of cDNA products were combined with 10 µL of TD buffer and 5 µL of ATM. Next, the mixture was incubated at 55C for 5 minutes, after which 5 µL of NT buffer was added, and the mixture was incubated at room temperature for 5 minutes. After this, the following components were added to the mixture to set up a PCR reaction: 8 µL of nuclease-free water, 1 µL of 10 µM P5_TSO primer, 1 µL of 10 µM P7_N70X_Custom2 primer, and 15 µL of Nextera PCR Master Mix. The thermal cycle program was: 95C 30s; 12 cycles of 95C 10s, 55C 30s, 72C 30s; 72C 5 min; 4C hold. Finally, the PCR products were cleaned up using 0.6X AMPure beads as above.

Library preparation of PLA products was performed as following. 1 µL of the pooled PLA products was used to set up a 20 µL PCR reaction, which contained 1X KAPA HiFi HotStart Readymix, 0.2 µM P5_TSO primer and 0.2 µM P7_N7XX_Custom2 primer. The thermal cycle program was: 95C 3 min; 12 cycles of 98C 20s, 67C 15s, 72C 20s; 72C 5 min; 4C hold. Then, the PCR products were cleaned up with 1.8X AMPure beads as above.

**Smart-Seq2 pipeline experiment**

96-well plates were first prepared by adding 4 µL of lysis buffer per well as per Smart-seq2 protocol[2]. After the PLA process, Jurkat and Raji cells were stained with propidium iodide (PI), and single, PI-negative cells were sorted into each well of two 96-well plates, one for each cell line. After sorting, the plates were centrifuged at 700g for at least 10s, and kept at -80C for storage. When ready to use, the plates were thawed on ice, incubated at 72C for 3 minutes, and centrifuged at 700g for at least 10s. Next, 2-4 µL of the cell lysate was used as input for PCR (the volume depends on how much lysate is needed to prepare the mRNA library, which can be skipped if one is only interested in the PLA aspect of Prox-seq). Each 25 µL PCR reaction also contained 1X KAPA HiFi HotStart Readymix, 0.1 µM Oligo_dTGT primer and 0.1 µM U_fwd primer. The thermal cycle program was: 98C 3 min; 5 cycles of 98C 20s, 55C 15s, 72C 1 min; 17 cycles of 98C 20s, 67C 15s, 72C 1 min; 72C 5 min; 4C hold.

After PCR, the products were cleaned up using 1.8X AMPure beads and Smart-seq2 protocol. Then, library preparation was performed to attach Illumina sequencing adapters, and to barcode the PLA products from each single cell with dual indices. To do this, 2 µL of PLA products from each well was used as input to a 25 µL PCR reaction, which contained 1X KAPA HiFi HotStart Readymix, 1 µM SmartPLA_P5_S5XX primer and 1 µM SmartPLA_P7_N7XX primer. The thermal cycle program was: 98C 3 min; 12 cycles of 98C 20s, 67C 15s, 72C 1 min; 72C 5 min; 4C hold.

After library preparation, the barcoded PLA products were cleaned up using 1.8X AMPure beads as above, and quantified with Qubit 1X dsDNA HS Assay Kit (Invitrogen, cat # Q33231). Then, the library from each well was normalized to 2 ng/ µL, and 2 µL from each well was pooled at equal volume into one 1.5 ml microfuge tube.

**Alignment of cDNA sequencing reads for Drop-seq pipeline**

First, the first 19 bases of read 2 were trimmed using the FASTX-toolkit (http://hannonlab.cshl.edu/fastx_toolkit/index.html) in order to remove the mosaic sequence from the Nextera transposase. Then, the read 1 and processed read 2 fastq files were used as input for Drop-seq tools v2.3.0 following the online instruction[1]. GRCh38 was used as the reference human genome. To extract the digital gene expression (DGE) matrix, we chose a list of cell barcodes according to the knee plot. Note that this list of cell barcodes will be later used for processing PLA sequencing reads.

**Alignment and processing of PLA sequencing reads**

To convert raw sequencing reads from PLA libraries to "digital PLA expression" (DPE) matrices, we developed a custom Java program that has two modes of processing, depending on whether the PLA libraries were prepared according to Drop-seq or Smart-seq2 pipeline. First, the program performs alignment by extracting the cell barcode, the UMI, and the PLA probe identities from the sequencing reads. For Drop-seq mode, the program will extract the cell barcode and UMI from read 1, and the sequence of the PLA product from read 2. (In this mode, the UMI on the PLA probe A is redundant and ignored.) Next, the program looks for valid PLA reads, which contain the connector region's sequence (TCGTGTCGTGTCGTGTCTAAAG) within 3 Levenshtein distance. We used the Levenshtein distance to allow for small insertion and deletion errors during PCR amplification and/or sequencing. Within the valid PLA reads, we look for the barcodes of antibodies A and B, and match them up against a text file containing the list of reference antibody barcodes and their targets. Any found antibody barcodes that are 1 Hamming distance or lower away from a reference antibody barcode are considered to be a match. After this, the alignment step is finished, and the output is a text file containing the cell barcode, the UMI, and the identities of probes A and B of each sequencing read. For Smart-seq2 mode, we extract the PLA sequence from read 1, and use the UMI on the PLA probe A. We do not need to extract the single-cell barcodes here, because they are automatically demultiplexed by Illumina's FastQ generation pipeline. Similar to the Drop-seq mode, we used the Levenshtein distance to identify valid PLA reads, and extract the identities of probes A and B. The output of this mode is multiple text files (one text file per single cell), each of which contains the UMI and the identities of probes A and B of each sequencing read.

After alignment, we perform cell barcode correction for Drop-seq libraries. This step is done to ensure that the barcodes obtained from PLA alignment match with the barcodes obtained from mRNA alignment. First, we import the list produced by Drop-seq tools v2.3.0 function *BamTagHistogram*, which contains the reference cell barcodes and their read counts. Then, we match the cell barcodes found from the PLA libraries to the reference list (a match has at most 1 Hamming distance). We do not perform cell barcode correction for Smart-seq2 libraries.

Next, we perform UMI merging for each single cell individually, by removing UMIs that are at most 1 Hamming distance away from another UMI that received more reads. This merging step is the same for both Drop-seq and Smart-seq2 libraries.

Lastly, we export the UMI merged reads into a DPE matrix as a text file. For Drop-seq mode, we choose a set of cell barcodes to be exported, either by making use of the "knee" plot for PLA reads, or by using the same set of cell barcodes chosen for Drop-seq tools v2.3.0 function *DigitalCount*. The output DPE matrix will have rows as different PLA products, and columns as different cell barcodes. For Smart-seq2 mode, we export all single cells in the DPE matrix, which has rows as different single cells, and columns as different PLA products.

**mRNA and PLA data analysis for Drop-seq pipeline**

The DGE matrix was processed using Seurat v3.0.2[3,4]. For pre-filtering, cells whose mitochondria genes' count exceeded 10% were removed, cells with fewer than 200 detected genes were removed, and genes detected in fewer than 5 cells were removed. Then, only the cells whose barcodes were present in both the filtered DGE matrix and the DPE matrix were retained.

The protein abundance was calculated from PLA data by calculating the number of UMI counts associated with each protein target, across both PLA probes A and B. For example, if PLA products CD3:CD28 and CD3:CD3 have a UMI count of 10 and 20, respectively, then the abundance of CD3 was $10\times1 + 20\times2 = 50$.

The data was then normalized (log-normalization for mRNA data, and centered log-ratio (CLR) transformation for protein abundance and PLA data), scaled, clustered (with resolution set to 0.3), and underwent dimensionality reduction with t-Distributed Stochastic Neighbor Embedding (t-SNE) visualization.

**PLA data analysis for Smart-seq2 pipeline**

The DPE matrix was processed with Python version 3.7.7 and RStudio. Cells with fewer than 500 UMIs were removed. A Python script was used to estimate the complex abundance using the UMI data. More details on this is available in another section below.

**Nearest neighbors aggregation**

The R package FNN was used to find the 5 nearest neighbors of each single cell using the mRNA data of the Drop-seq data set. The gene and protein's normalized expression level of each single cell was then calculated from the average of the cell's nearest neighbors.

**Expected PLA count**

We reason that, in the case where no complexes are present, the expected count of a PLA product i:j, Ei,j, in each single cell is equal to the product of the marginal probability of probe A targeting protein i, the marginal probability of probe B targeting protein j, and the total PLA count:

$$E_{i,j} = \frac{\sum_{k=1}^{n} X_{i,k} \times \sum_{k=1}^{n} X_{k,j}}{\sum_{k=1}^{n} \sum_{k=1}^{n} X_{k,l}}$$

(1)

If Xi,j is different from Ei,j for any PLA product i:j, then at least one protein complex exists in the data set. Indeed, our simulated PLA data support this (Supplementary Fig. 6).

**Linear regression and likelihood ratio test**

Another way to test whether PLA data contains protein complex information is to perform linear regression. We modeled the observed count of PLA product i:j using the null model and the alternative model:

$$\mathrm{H_0}: X_{i,j} \sim \beta_0 + \beta_1 \sum_{k=1}^{n} X_{i,k} + \beta_2 \sum_{k=1}^{n} X_{k,j}$$

(2)

$$\mathrm{H_a}: X_{i,j} \sim \beta_0 + \beta_1 \sum_{k=1}^{n} X_{i,k} + \beta_2 \sum_{k=1}^{n} X_{k,j} + \beta_\mathrm{x} \left( \sum_{k=1}^{n} X_{i,k} \times \sum_{k=1}^{n} X_{k,j} \right)^{1/2}$$

(3)

Thus, β1 and β2 represented the effect size of the abundance of detected probes Ai and Bj, respectively, and βx represented the effect size of the interaction between abundance of probes Ai and Bj. The interaction term was square rooted so that the magnitude of βx was comparable to that of β1 and β2. We then used the likelihood ratio test to see if the alternative model $H_a$ was significantly better than the null model $H_0$ in explaining the data. If it was for the majority of PLA products, then the interaction effect was present, suggesting that the observed PLA data contained protein complex information.

Linear regression and likelihood ratio test was performed using RStudio.

**Simulated PLA data**

To help with developing a protein complex calling method, we wrote a Python script to simulate PLA data. Let n denote the number of protein targets. For each single cell, there will be $n^2$ possible PLA products. Let $X_{i,j}$ denote the count of PLA product between probes Ai and Bj, which target protein i and j, respectively (i and j are from 1 to n). Xi,j depends on the abundance of protein complex i:j (if it exists, Supplementary Fig. 5a), and the ligation of Ai and Bj that are in proximity to each other by chance (Supplementary Fig. 5b,c). Assuming that non-specific binding of antibodies is minimal, then Xi,j depends on the abundance of complex i:j, and the abundance of protein i and protein j. We further assume that all protein molecules are bound by their corresponding probes, which means that we ignore any differences in antibody binding.

For each single cell, we can simulate proximity ligation by ligation of pairs of probes that are randomly distributed on a sphere: any pairs of probe A and B within the ligation distance can produce a PLA product. Random generation of probe A and B is achieved in two steps. First, we generate the complexes as random points on a sphere of 10,000 units in diameter (one unit is equal to 1 nm). These complexes are represented as an input n-by-n matrix **A**, where the element $a_{i,j}$ on row i and column j is equal to the abundance of complex i:j (if the complex i:j does not exist, $a_{i,j}$=0). Because of the assumptions above, each complex i:j location contains a probe Ai and a probe Bj.

Second, we generate the protein molecules that do not form a complex with any other probe targets, also as random points on the same sphere. Biologically, these proteins can be in monomer forms, or they form complexes with proteins that are not part of the current antibody panel. These proteins are represented by two n-by-1 input vectors **x** and **y**, where the element $x_i$

denotes the number of protein i molecules bound by probe $A_i$, and element $y_i$ denotes the number of protein i molecules bound by probe $B_i$.

Finally, we calculate the pairwise Euclidean distance between all probes A and all probes B. Any probe Ai-probe Bj pair with a distance less than or equal to the ligation distance (assumed to be 50 units) produces one PLA product i:j.

We also introduce single cell heterogeneity in our simulation by including a cell-wide scaling factor for the amount of probes (the factor is log-normal distributed with $\mu = 0$ and $\sigma = 0.5$).

**Estimation of complex abundance**

Using the expected count from Eq. (1), one would naturally expect that for a complex i:j, the observed count Xi,j must be higher than the expected count Ei,j. However, as shown in the example in Supplementary Fig. 5, both products 1:1 and 2:2 display this feature, even though only 2:2 is a complex.

The most unique feature of our assay is that it enables combinatorial screening of protein complexes and protein interactions. We reason that maximizing the true positive rate is more important than minimizing the false discovery rate. As a result, we aim to find candidate complexes, which are the PLA products whose observed counts are higher than expected counts (eg, 1:1 and 2:2 in Supplementary Fig. 6). As discussed in the previous section, the observed count is assumed to be the sum of two components, the abundance of the complexes and the random ligation of proximal probe pairs. It follows that, if we subtract the complex abundance from the observed PLA count, the remaining count represents the random ligation component, which can be estimated using Eq. 1. Thus, the abundance of a complex i:j, $D_{i,j}$, satisfies the following equation:

$$X_{i,j} - D_{i,j} = \frac{\left(\sum_{k=1}^{n} X_{i,k} - \sum_{k=1}^{n} D_{i,k}\right) \times \left(\sum_{k=1}^{n} X_{k,j} - \sum_{k=1}^{n} D_{k,j}\right)}{\sum_{k=1}^{n}\sum_{l=1}^{n} X_{k,l} - \sum_{k=1}^{n}\sum_{l=1}^{n} D_{k,l}}$$

(4)

The left hand side of the equation is the random ligation component of PLA product i:j (in other words, the adjusted PLA count). The right hand side is the expected count for i:j calculated using adjusted PLA counts. Di,j must be non-negative, and if Di,j = 0, then i:j is not a true, existing protein complex. Solving Di,j for all PLA products involves solving a system of $n^2$ quadratic equations, which can have multiple sets of solutions. We can arrive at a set of solutions using an iterative algorithm:

$$D_{i,j}^{(m+1)} = X_{i,j} - \frac{\left(\sum_{k=1}^{n} X_{i,k} - \sum_{k=1}^{n} D_{i,k}^{(m)}\right) \times \left(\sum_{k=1}^{n} X_{k,j} - \sum_{k=1}^{n} D_{k,j}^{(m)}\right)}{\sum_{k=1}^{n}\sum_{l=1}^{n} X_{k,l} - \sum_{k=1}^{n}\sum_{l=1}^{n} D_{k,l}^{(m)}}$$

(5)

where $D_{i,j}^{(m)}$ is the estimated complex abundance of i:j at the m$^{th}$ iteration, and $D_{i,j}^{(0)} = 0$ for all i,j. The iterative process is stopped either when the solutions converge, or when the maximum number of iterations is reached. Using the same simulated data set from Supplementary Fig. 6b, we can see that the estimated complex abundance correlates with the true complex abundance (Supplementary Fig. 10a).

To ensure that Di,j is non-negative, during each iteration m+1, we performed a one-tailed one sample t-test on $D_{i,j}^{(m+1)}$ of all single cells and a cutoff value. If the null hypothesis that the mean of $D_{i,j}^{(m+1)}$ was lower than the cutoff value was not rejected, we set $D_{i,j}^{(m+1)} = 0$ for all single cells. We chose the cutoff value in this study to be 1, because we observed that a cutoff value of 1 resulted in more isotype control PLA products being rejected than a cutoff value of 0.

We also added another option to enforce symmetry: that if i:j was detected to be a complex, then j:i should also be detected as a complex. To do this, we introduced a parameter called sym_weight, such that if $D_{i,j}^{(m+1)} > 0$ but $D_{j,i}^{(m+1)} = 0$, then the latter was recalculated as $D_{i,j}^{(m+1)} = \text{sym\_weight} \times D_{i,j}^{(m+1)}$. In this study, we arbitrarily chose sym_weight=0.25. We note that for our dataset, this parameter only led to a few symmetrical PLA products being detected as complexes, and not in any fundamentally new complexes.

As expected, the iterative algorithm also misidentifies 1:1 as a complex (Supplementary Fig. 10b). This also causes the estimate abundance of 2:2 to be lower than the true abundance. After subtracting the complex signal component, we observe that the adjusted observed PLA counts agree with the expected counts (Supplementary Fig. 10c). This is because the system of quadratic equations in Eq. 2 has multiple sets of solutions, and the iterative algorithm did not arrive at the true solution. We do note that, if we had a priori knowledge that 1:1 does not exist as a homodimer, we could force $D_{1,1}$ to be zero for all iterations. Then, we can arrive at a set of solutions much closer to the true solutions (Supplementary Fig. 10d). At the moment, we do not have a method that allows us to distinguish false positive complexes from true positive ones. Being able to constraint the false positives will enable the iterative algorithm to estimate the complex abundance more accurately, and to identify low abundance complexes.

1. Macosko, E. Z. *et al.* Highly parallel genome-wide expression profiling of individual cells using nanoliter droplets. *Cell* **161**, 1202–1214 (2015).

2. Picelli, S. *et al.* Full-length RNA-seq from single cells using Smart-seq2. *Nat. Protoc.* **9**, 171–181 (2014).

3. Butler, A., Hoffman, P., Smibert, P., Papalexi, E. & Satija, R. Integrating single-cell transcriptomic data across different conditions, technologies, and species. *Nat. Biotechnol.* **36**, 411–420 (2018).

4. Stuart, T. *et al.* Comprehensive Integration of Single-Cell Data. *Cell* **177**, 1888-1902.e21 (2019).
